# Ibogaine is associated with reorganization of high-beta brain networks in veterans with post-traumatic stress disorder

**DOI:** 10.64898/2026.03.20.713241

**Authors:** Kenneth Shinozuka, Mattia Rosso, Anna Chaiken, Jennifer I. Lissemore, Reece Jones, Nelson Descalco, Venkatesh Subramani, Maarten Belgers, Kirsten N. Cherian, Martijn Arns, Davide Momi, Raag D. Airan, Leonardo Bonetti, Arnt Schellekens, Maheen M. Adamson, Corey J. Keller, Cammie Rolle

## Abstract

Post-traumatic stress disorder (PTSD) is a debilitating condition that affects millions of veterans. A recent observational study in 30 veterans showed that a single dose of the atypical psychedelic ibogaine can be highly effective at treating PTSD up to twelve months later. Although a prior study demonstrated that ibogaine transiently alters electroencephalography (EEG) power in various frequency bands, the long-term, network-level neural mechanisms targeted by ibogaine are unclear. Here, we investigated whether ibogaine-related clinical improvements are associated with the reorganization of certain brain networks. We applied a novel framework, FREQuency-resolved brain Network Estimation via Source Separation (FREQ-NESS), to identify frequency-specific brain networks in resting-state EEG data acquired at baseline, three to four days after treatment (immediate-post), and one month after ibogaine. At both the immediate-post and one month-post timepoints, high-beta (24 and 25 Hz) networks shifted away from frontal areas and towards posterior regions, an effect that was replicated in an independent EEG dataset on ibogaine treatment for opioid use disorder. This posterior shift was significantly correlated with improvements in PTSD symptoms at both timepoints. Neural field modeling demonstrated that these posterior high-beta shifts are associated with increases in corticocortical, but not corticothalamic, connectivity. Our results are consistent with prior evidence implicating aberrant frontal beta-band activity in PTSD. Overall, we demonstrate that the reconfiguration of high-beta brain networks could be a robust biomarker for ibogaine’s therapeutic effects.

## Section 1. Introduction

Psychedelic drugs have recently attracted renewed interest due to growing evidence that they can treat a wide range of psychiatric conditions, including post-traumatic stress disorder (PTSD), depression, anxiety, and substance use disorders (Beutler et al., 2024; Cherian, Shinozuka, et al., 2024; Evans et al., 2024; Muir et al., 2024; Shinozuka, Tabaac, et al., 2024b, 2024a; Tabaac et al., 2024a, 2024b). One of these psychedelics is ibogaine, the primary psychoactive molecule in *Tabernanthe iboga*, a shrub from Gabon and the Congo that has been ingested as a spiritual sacrament by the Bwiti religion for millennia (Ravalec et al., 2007). Unlike “classic” psychedelics such as lysergic acid diethylamide (LSD) and psilocybin, ibogaine is considered to be an “atypical” psychedelic because its psychoactive effects do not appear to be mediated by the 5-HT_2A_ receptor, but instead may involve the kappa-opioid and N-methyl-D-aspartate receptor (Cherian, Shinozuka, et al., 2024; Rodrıguez et al., 2020).

Using the Magnesium-Ibogaine: the Stanford Traumatic Injury to the CNS (MISTIC) protocol, a recent observational study in 30 U.S. veterans demonstrated that ibogaine was associated with marked improvements in symptoms related to traumatic brain injury (TBI). Reductions in PTSD, depression, anxiety, and functional disability exhibited very large effect sizes; for instance, the Cohen’s *d* for improvements in PTSD symptoms one month after ibogaine was 2.54 (Cherian, Keynan, et al., 2024). These effects were sustained at least 12 months after ibogaine treatment (Faerman, Lissemore, et al., in press). Preliminary evidence also indicates that ibogaine could be highly effective at treating substance use disorders, including opioid and cocaine addiction (Cherian, Shinozuka, et al., 2024; Keighley et al., 2025; Mash et al., 2018; Nicolas, 2025; Noller et al., 2018; Prior & Prior, 2014).

Yet the neural mechanisms underlying the long-term therapeutic effects of ibogaine remain unclear. EEG recordings of the same veterans demonstrated increases in lower-frequency power (e.g., theta, alpha) and decreases in higher-frequency power (e.g., beta, gamma) three to four days after ibogaine treatment, but these effects tended not to be sustained one month later (Lissemore, Chaiken, et al., 2025). Decreases in peak alpha frequency were correlated with PTSD symptom improvements at one month, yet such decreases also occur during aging and cognitive impairment (Garcés et al., 2013; Park et al., 2024; Zhao et al., 2025). Meanwhile, ibogaine appears to reduce the estimated age of the brain, based on machine learning analyses of structural MRI data (Geoly et al., 2026), while improving multiple domains of cognition (Cherian, Keynan, et al., 2024).

Conventional spectral approaches, such as averaged power and phase-based functional connectivity, only measure neural activity in individual regions or pairs of regions, rather than capturing the coordination of oscillatory dynamics across distributed, whole-brain networks.

However, in PTSD, converging evidence suggests that symptoms arise not simply from nonspecific increases or decreases in power, but from maladaptive patterns of network organization in which default mode, control, and salience systems interact inefficiently (Akiki et al., 2017; Lanius et al., 2015; Ross & Cisler, 2020). As a result, approaches that focus exclusively on spectral amplitude or pairwise connectivity may overlook clinically meaningful reconfigurations of network architecture that are central to the persistence and resolution of PTSD symptoms.

A novel framework, FREQuency-resolved Network Estimation via Source Separation (FREQ-NESS), was recently developed to measure these network-based changes (Rosso et al., 2025). This method identifies brain networks that oscillate at specific frequencies, by determining the weights of brain regions that maximally separate narrow-band covariance (i.e., covariance when the data is narrowly filtered to the frequency of interest) from broadband covariance. FREQ-NESS reveals the brain networks that best explain the connectivity at a particular frequency. This approach has several advantages over traditional methods for measuring functional connectivity in EEG data. First, it measures coordinated whole-brain network patterns, rather than connectivity between individual pairs of regions. Second, and most importantly, it isolates oscillatory networks that are driven by frequency-specific covariance rather than broadband, spatially diffuse fluctuations in power, yielding a more robust and interpretable characterization of frequency-resolved brain organization. Finally, because FREQ-NESS identifies changes in brain networks at individual frequencies, it can reveal frequency-specific network reorganization outside canonical bands, underscoring its utility for discovery-driven analyses. For instance, the method was recently used to show that LSD shifts high-alpha (12-13 Hz) and low-alpha (8 Hz) networks in opposite directions, towards posterior and towards anterior cortex, respectively (Shinozuka, Rosso, et al., 2025). In summary, FREQ-NESS is well-suited for detecting large-scale network reorganization following interventions such as psychedelics.

While FREQ-NESS characterizes which networks change, neural field models provide a clear biophysical framework for identifying the neural mechanisms of frequency-specific network reorganization. This class of models describes the spatiotemporal evolution of population-level neural activity by treating the cortex as a continuous sheet where activity propagates via spatially distributed connectivity, resulting in large-scale oscillatory dynamics (Sanz-Leon et al., 2018). The Robinson-Rennie-Wright (RRW) model extends neural field theory by describing how cortico-cortical, cortico-thalamic, and intrathalamic connectivity jointly generate canonical EEG rhythms such as alpha and beta (Abeysuriya et al., 2014; Abeysuriya & Robinson, 2016; Robinson et al., 1997, 2003). RRW is well-suited for modeling EEG data on psychedelics due to the popular theory that psychedelics dramatically increase the vividness of sensory perception by elevating connectivity between the cortex and the thalamus, a brain region known for gating, integrating, or inhibiting irrelevant sensory stimuli (Avram et al., 2021; Coleman et al., 2025; Müller et al., 2017; Onofrj et al., 2023; Vollenweider & Geyer, 2001). RRW can therefore be used to assess how ibogaine reshapes network topographies by altering cortico-thalamic connectivity.

Here, we investigated whether the therapeutic effects of ibogaine are associated with the reorganization of frequency-specific brain networks, by applying FREQ-NESS to resting-state EEG data acquired before, immediately after, and one month after ibogaine administration (Cherian, Keynan, et al., 2024). We also replicated our results in an independent sample of patients who were treated with ibogaine for opioid use disorder. We hypothesized that ibogaine reshapes the spatial organization of frequency-specific brain networks by altering cortico-thalamic coupling, as modeled by RRW, leading to sustained network reconfiguration that correlates with improvements in PTSD symptoms.

## Section 2. Methods

### Section 2.1. Participants, treatment, and clinical measures

The data were acquired as part of the previous MISTIC study (Cherian, Keynan, et al., 2024). Details are summarized in **Supplementary Methods 1-3.** In brief, 30 male veterans with mild-to-moderate TBI received a single dose of ibogaine (12.1 ± 1.2 mg/kg) and were pre-treated with intravenous magnesium sulfate (1 g) to mitigate cardiovascular risks, particularly Q-T interval prolongation. 23 of the 30 participants were diagnosed with PTSD based on the Mini International Neuropsychiatric Interview. Using the Clinician-Administered PTSD Scale for DSM-5 (CAPS-5), PTSD symptoms were measured at baseline, immediate-post (3-4 days), and one month-post ibogaine.

### Section 2.2. EEG data acquisition and preprocessing

EEG data were collected at three timepoints: baseline (2-3 days before ibogaine treatment; *n* = 30), immediate-post (3-4 days after ibogaine treatment; *n* = 30), and one month-post ibogaine (*n* = 27; three participants were not recorded with EEG due to logistical issues). EEG data were acquired and preprocessed as described previously (Lissemore, Chaiken, et al., 2025) and as summarized in **Supplementary Methods 4-5.**

### Section 2.3. FREQ-NESS

As described previously (Rosso et al., 2025; Shinozuka, Rosso, et al., 2025), FREQ-NESS applies generalized eigendecomposition (GED) to extract brain networks that operate at specific frequencies (M. X. Cohen, 2017, 2022; Rosso et al., 2021, 2022, 2023). We performed FREQ-NESS on the sensor-space EEG data (**Figure 1**). For each target frequency, GED computes electrode weights that best distinguish the covariance of the signal narrowly filtered around that frequency from that of the broadband signal 𝐗_𝐛𝐫𝐨𝐚𝐝_(64 x *T*, where 64 is the number of channels and *T* is the number of timepoints) thereby determining frequency-specific network patterns. Mathematically, this is achieved by solving the equation:

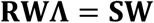

**Figure 1.**
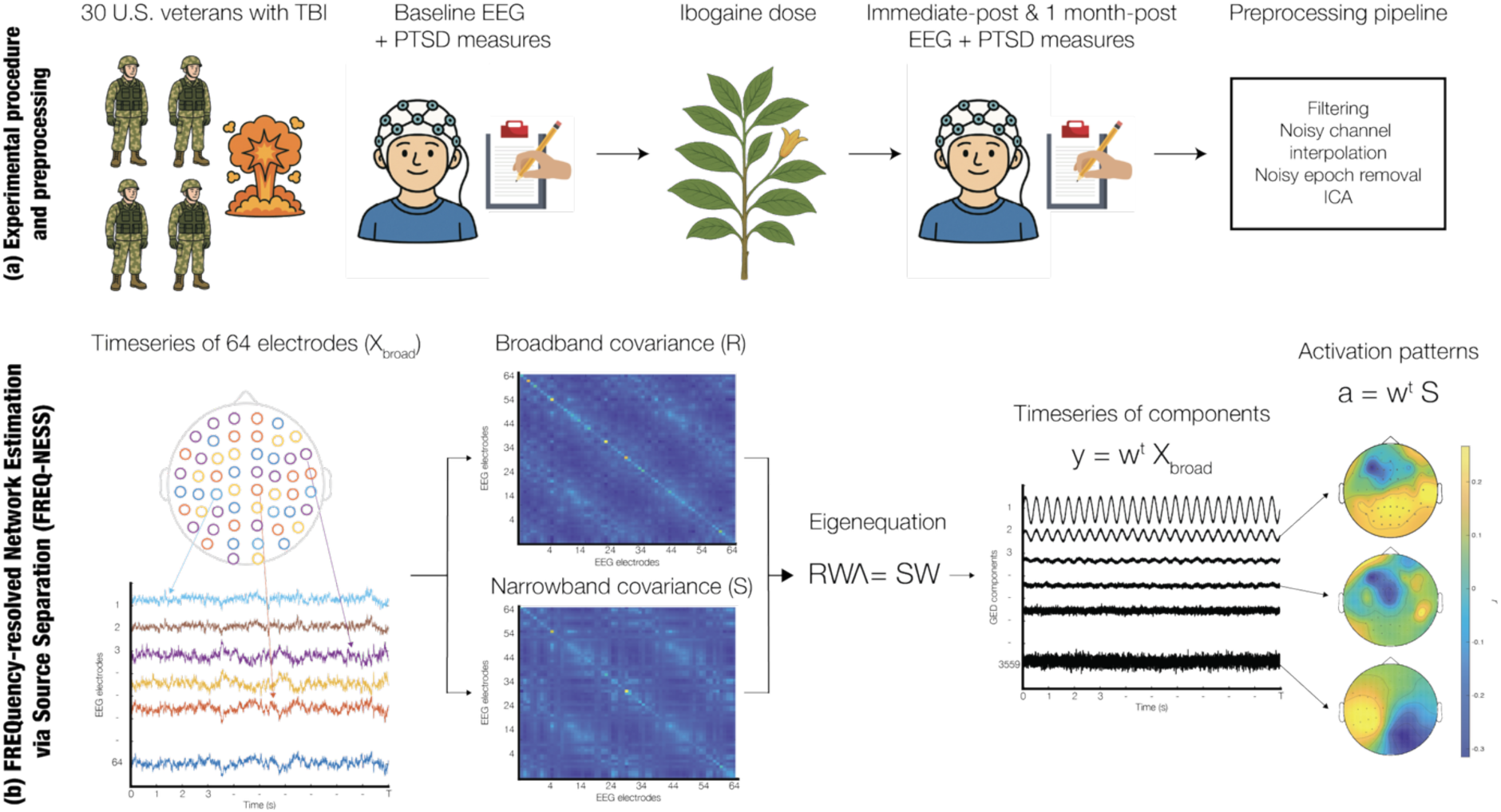
Overview. (a) A single dose of ibogaine (mean total dose = 12.1 ± 1.2 mg per kg) was administered to 30 U.S. Special Operations Forces veterans with mild-to-moderate traumatic brain injury. EEG and PTSD severity were measured at baseline, immediate-post (3-4 days after), and 1 month-post ibogaine. EEG preprocessing steps included filtering, noisy channel interpolation, noisy epoch removal, and independent component analysis (ICA). (b) The 64-channel timeseries are narrow-band filtered at a range of 29 equally spaced frequencies (2-30 Hz). For each frequency, the narrowband covariance matrix, **S**, is computed. The sets of channel weights **W** that maximally separate **S** from the broadband covariance matrix, **R**, are determined by solving the eigenequation **RWΛ** = **SW**, where **Λ** contains the eigenvalues or variance explained by each set of channel weights. **W** is projected onto electrode space to obtain the network activation patterns, or spatial topographies, **a**. We measured the effect of ibogaine on the network **a** that was associated with the highest eigenvalue in **Λ**.

where 𝐖 is a 64 × 64 matrix of eigenvectors representing electrode weights, 𝚲 is a diagonal matrix of eigenvalues, 𝐑 is the broadband covariance matrix, and 𝐒 is the narrow-band covariance matrix at the frequency of interest. Each eigenvalue reflects the relative expression (i.e., ratio) of narrow-band covariance to broadband covariance captured by the corresponding eigenvector, with larger eigenvalues indicating network patterns that are more strongly expressed at the frequency of interest compared to broadband activity.

To obtain network timeseries, each eigenvector 𝐰 is projected into electrode space as:

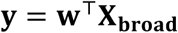

Likewise, to obtain spatially interpretable network maps, each eigenvector 𝐰 is projected into electrode space as:

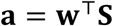

producing network activation patterns 𝐚 (64 x 1) that represent the spatial distribution of network activity (Haufe et al., 2014).

The EEG signal was narrow-band filtered using Gaussian wavelets with a 0.3 Hz half-maximum width centered on each frequency, based on previous applications of GED to EEG data (Rosso et al., 2021, 2023). Twenty-nine frequencies between 2 Hz and 30 Hz were analyzed, spaced apart by intervals of 1 Hz. These frequencies span the delta to beta bands, avoiding gamma since it tends to be confounded by muscle artifacts (Muthukumaraswamy, 2013), and avoiding arbitrary band limits, such as 8-13 Hz for alpha. For a complete mathematical description of FREQ-NESS, see Rosso et al. (2025).

Linear mixed-effects models (LMEs) were used to measure statistically significant changes in the leading eigenvalue per frequency, and cluster-based permutation testing was used to assess statistically significant changes in network topographies per frequency. Only the leading eigenvalue/eigenvector was tested statistically to reduce multiple comparisons. In each case, the false discovery rate (FDR) was used to correct for multiple comparisons across frequencies and timepoints. Further details are provided in **Supplementary Methods 6.**

FREQ-NESS was also compared to traditional methods for measuring functional connectivity, namely the weighted phase lag index (wPLI), as described further in **Supplementary Methods 7**.

To ensure that our results were not confounded by any muscle or ocular artifacts, we ran three control analyses, as discussed further in **Supplementary Methods 8.**

### Section 2.4. Correlations between network topographies and PTSD improvements

For networks that exhibited significant topographical changes under ibogaine, Spearman correlations were computed between the network activation at each electrode and total CAPS-5 scores, resulting in a CAPS-5 “correlation map.” Further Pearson correlations were computed between this correlation map and the *y*-coordinate of each electrode, reflecting the position of the electrode along the anterior-posterior axis of the brain. The same procedure was repeated for CAPS-5 subscales B, C, D, and E, which together constitute the core symptoms of PTSD. To correct for multiple comparisons, *p*-values of the Pearson correlations were adjusted with FDR. There was a total of 20 comparisons (2 networks with significant changes in network topography [24 and 25 Hz] x 2 study timepoints [immediate- and one month-post] x 5 scores, including total CAPS-5 score and four subscale scores).

To test whether the anterior-posterior topographic gradient of the correlation between network activation change and CAPS-5 scores differed across symptom subscales and frequency bands, we fit an LME. The dependent variable was the electrode-wise Pearson correlation coefficient between the change in network activation and CAPS-5 subscale score . Fixed effects included electrode anterior-posterior position (y-coordinate, continuous), CAPS-5 subscale (Intrusion [reference], Avoidance, Cognition/Mood, Arousal/Reactivity), frequency band (24 Hz [reference], 25 Hz), and all two- and three-way interactions. A random intercept for electrode was included to account for repeated observations within each electrode across conditions. The three-way interaction (y-coordinate x subscale x frequency) was evaluated via both a likelihood ratio test comparing the full model to a reduced model omitting the three-way interaction terms and a Type III ANOVA with Satterthwaite’s method.

### Section 2.5. Neural field modeling

To explain the changes in network topographies based on corticothalamic connectivity, we employed the Robinson-Rennie-Wright (RRW) corticothalamic neural field model, implemented in the BrainTrak software suite (Abeysuriya & Robinson, 2016). The model was first fit to the EEG power spectra, and then model parameters were used to simulate high-beta network topographies (see **Supplementary Methods 9**). The RRW model represents large-scale brain dynamics using four spatially distributed neural populations: cortical excitatory (*e*) and inhibitory (*i*) neurons, thalamic relay (*s*) neurons, and thalamic reticular (*r*) neurons. These populations interact through excitatory and inhibitory connections characterized by loop gains *G*_ee_, *G*_ei_, *G*_ese_, *G*_esre_, and *G*_srs_, which together define the strength of cortico-cortical, cortico-thalamic, and intrathalamic feedback pathways. Each population’s mean soma potential evolves according to a second-order differential operator that models synaptodendritic filtering, while axonal propagation is represented by a damped wave equation incorporating finite conduction velocity and propagation range (see *Section 2.11*). The core dynamics of the model are captured by three parameters:

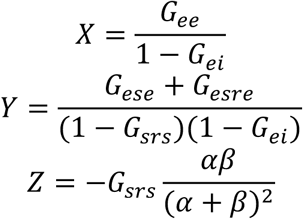

where 𝛼 and 𝛽 are the decay and rise rate of cell-body potential, respectively.

The model is linearized about its steady state (a “baseline” in which the neural dynamics do not change) to yield an analytic transfer function describing how neuronal population activity generates the EEG power spectrum. The BrainTrak framework fits this theoretical spectrum to empirical EEG spectra using a Markov Chain Monte Carlo (MCMC) algorithm. Physiologically plausible priors constrain each parameter within known neurobiological ranges. Fitting yields posterior distributions for the loop gains and consequently *X*, *Y*, and *Z*.

In this study, all of the free parameters, which include the loop gains, 𝛼, 𝛽, and the corticothalamic loop delay, were fitted to the baseline EEG power spectra. 𝛼, 𝛽, and the corticothalamic loop delay were kept fixed when fitting the immediate-post and 1 month-post power spectra. (Thus, there were eight free parameters at baseline, and five free parameters at the other two timepoints.) *X*, *Y*, and *Z* were calculated based on the loop gains. Since Braintrak fits parameters to each electrode, we computed the median of each parameter across electrodes, weighted by the inverse goodness of fit (χ^2^), for each subject. To assess significant changes in the weighted median of *X*, *Y*, and *Z*, we used an LME with timepoint as a categorical fixed effect and subject-specific random intercepts.

High-beta network topographies were then simulated based on the fitted parameters, as described further in **Supplementary Methods 9.**

### Section 2.6. Replication of FREQ-NESS results in an independent dataset

We sought to replicate our main result in an independent dataset, in which 14 patients with opioid use disorder (OUD) were administered a single 10 mg/kg dose of ibogaine in an open-label fashion (Knuijver et al., 2022). No magnesium pre-treatment occurred in this study. Usable resting-state EEG was obtained from 11 out of 14 participants at baseline and a median of 12 days (± 8.15 days) after ibogaine. Further details on the EEG acquisition are provided in **Supplementary Methods 10.** Preprocessing of this EEG data was similar to that of the MISTIC dataset. One participant who exhibited extreme outliers in high-beta network activation (based on a *z*-score threshold of |*z*| > 3) was removed from the analysis.

## Section 3. Results

### Section 3.1. Ibogaine is associated with posterior shifts in high-beta networks

FREQ-NESS was used to identify brain networks oscillating at specific frequencies ranging from 2 to 30 Hz. While ibogaine did not significantly change the overall prominence (i.e., variance explained) of frequency-specific brain networks, it did significantly alter the spatial organization of high-beta (24–25 Hz) networks immediately after treatment and at one month follow-up (**Figure 2a-d**). Ibogaine significantly reduced the activation of frontal left electrodes in these networks, while significantly elevating the activation of posterior electrodes, especially at the one month-post timepoint (24 Hz, immediate-post > baseline: cluster-level *p*_FDR_ = 0.0039; 1 month-post > baseline: cluster-level *p*_FDR_ = 0.0039; 25 Hz, immediate-post > baseline: cluster-level *p*_FDR_ = 0.0116; 25 Hz, 1 month-post > baseline: cluster-level *p*_FDR_ = 0.0116). (Further cluster statistics are reported in **Table 1**.) This “posterior shift” was significantly greater in the high-beta frequencies than in other frequency bands; for example, the 5 Hz network exhibited no such shift at either timepoint (**Figure 2e-f**). The high-beta activation of a representative frontal left electrode (F5) was significantly lower than that of delta, theta, alpha, or low-beta activation (**Figure 2g**). Likewise, the high-beta activation of a representative posterior electrode (Pz) was significantly greater than that of the other frequency bands (**Figure 2h**). The high-beta posterior shift also survived a sensitivity analysis in which one outlier was removed, while also becoming significant in other high-beta networks, such as 23 Hz (**Supplementary Results 2**; **Supplementary Figure 2**).

**Figure 2.**
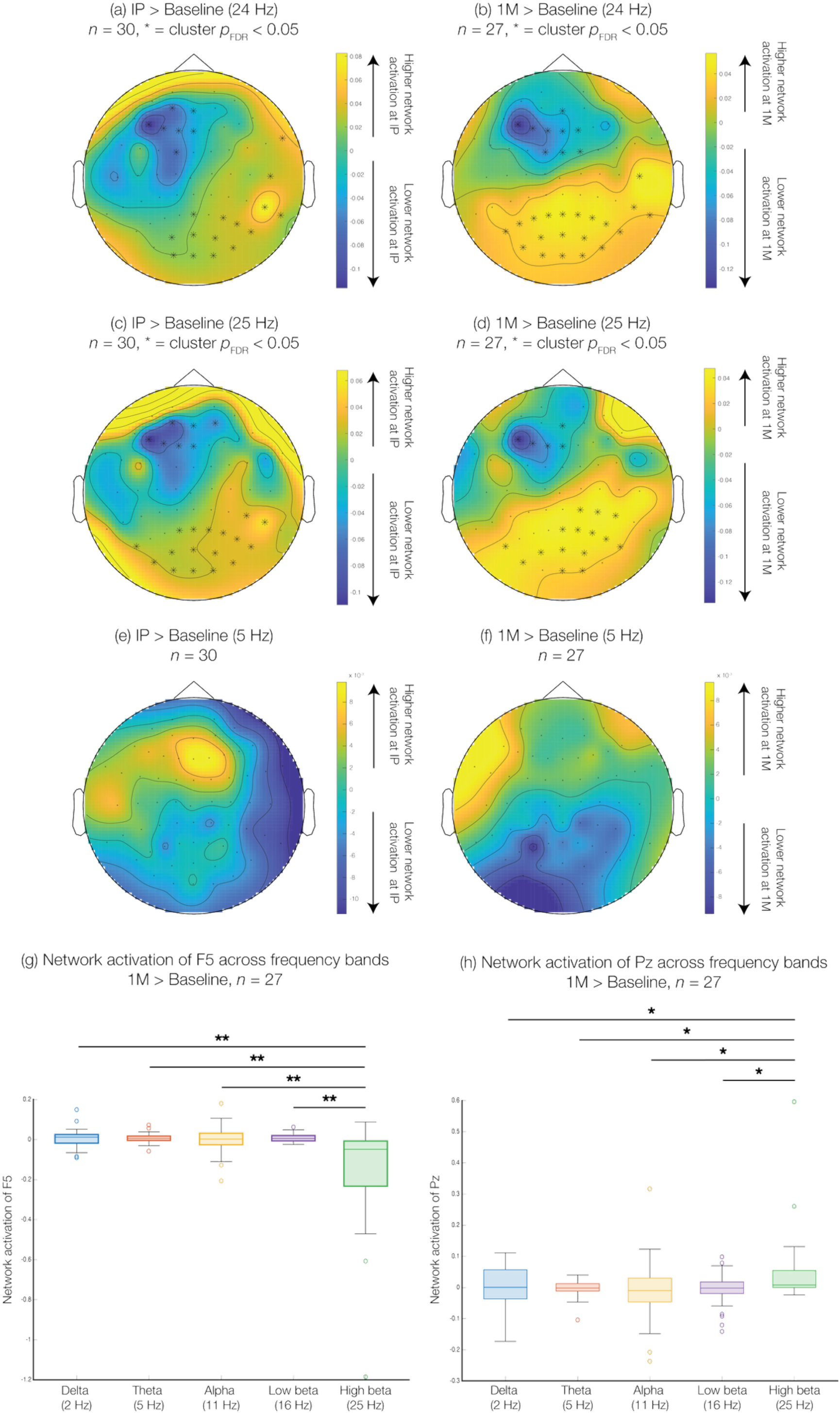
Ibogaine treatment is associated with posterior shifts in high-beta (24, 25 Hz) network topographies. Cluster-based permutation testing revealed significant changes in network topographies at 24 Hz, from baseline to both of the other timepoints (a, b), and at 25 Hz, also from baseline to both of the other timepoints (c, d). Brighter colors indicate increased activation of electrodes within the corresponding network, relative to baseline, while darker colors denote decreased activation relative to baseline. Asterisks indicate significant electrodes. In all cases, ibogaine significantly elevated the activation of posterior electrodes while reducing the activation of left frontal electrodes. (e, f) Ibogaine did not have a significant effect on the network topographies of any other frequencies, such as 5 Hz. (g) Ibogaine was associated with a significant decrease in the 25 Hz network activation at a representative left frontal electrode, relative to the network activation of frequencies in other bands. (h) Conversely, after ibogaine, there was a significant increase in the 25 Hz network activation at a representative posterior electrode, relative to the network activation of frequencies in other bands. IP = immediate-post; 1M = 1 month-post.

**Table 1.** Statistics of significant clusters of changes in high-beta network topographies. Cluster size, channel where the test statistic (*t*-value) reaches its absolute maximum and the value of that statistic, cluster-level *p*-value, and FDR-adjusted cluster-level *p*-value. For the latter, FDR was applied to a total of 58 comparisons (29 frequencies x 2 planned timepoint contrasts, i.e., immediate-post vs. baseline and one month-post vs. baseline). *n* = 27 participants were included in the analysis to account for three participants who did not have data at the one month-post timepoint.

| | Cluster Size | Max $t$ | Cluster $p$ | FDR-adjusted $p$ |
| --- | --- | --- | --- | --- |
| 24 Hz,<br>Immediate-post<br>> Baseline | 15 | POz: $t = 4.198$ | $p = 0.0002$ | $p_{\text{corrected}} = 0.0039$ |
| 24 Hz,<br>1 month-post<br>> Baseline | 20 | Oz: $t = 3.160$ | $p = 0.0002$ | $p_{\text{corrected}} = 0.0039$ |
| 25 Hz,<br>Immediate-post<br>> Baseline | 17 | POz: $t = 3.574$ | $p = 0.0008$ | $p_{\text{corrected}} = 0.0116$ |
| 25 Hz,<br>1 month-post<br>> Baseline | 15 | P6: $t = 2.499$ | $p = 0.0002$ | $p_{\text{corrected}} = 0.0116$ |

We also performed several control analyses to ensure that the posterior shifts could not be attributed to muscle or ocular artifacts (**Supplementary Figure 3**). A proxy of electromyography (EMG) activity – mean frontal and temporal power in the 40-80 Hz band – was not significantly associated with posterior shifts (24 Hz, immediate-post > baseline: *p* = 0.50; 1 month-post > baseline: *p* = 0.10; 25 Hz, immediate-post > baseline: *p* = 0.69, 1 month-post > baseline: *p* = 0.14). The posterior shifts were also not affected by excluding epochs with high-amplitude electrooculography (EOG) proxies or re-referencing the data (24 Hz, immediate-post > baseline: *p* = 0.28; 1 month-post > baseline: *p* = 0.28; 25 Hz, immediate-post > baseline: *p* = 0.69; 1 month-post > baseline: *p* = 0.41). The observed changes in high-beta network topographies are therefore robust to common EEG artifacts.

A more conventional EEG-based measure of functional connectivity, i.e., wPLI, was unable to detect an effect of ibogaine on high-beta brain networks. Unlike FREQ-NESS, which measures whole-brain patterns of covariance, wPLI captures connectivity between pairs of electrodes. After multiple comparisons were corrected with the network-based statistic, ibogaine did not have a significant effect on wPLI when the EEG signal was narrowband-filtered to either 24 or 25 Hz. There were no significant differences in wPLI across the whole alpha (8-13 Hz) or beta (13-30 Hz) bands either. Further results are discussed in **Supplementary Results 3** (**Supplementary Figure 4**).

### Section 3.2. The posterior shift is replicated in an independent dataset on ibogaine

The posterior high-beta network shift computed by FREQ-NESS was also observed in an independent resting-state EEG dataset of patients with OUD who were administered a single dose of ibogaine (**Figure 3**). Even though several frontal and occipital electrodes were either not recorded or removed from analysis in this dataset, both 24 and 25 Hz network activation nevertheless increased in posterior electrodes while decreasing in anterior electrodes, especially left frontal electrodes, after ibogaine treatment. Clusters exhibiting these changes were statistically significant in the 25 Hz network (cluster-level *p*_FDR_ = 0.03; further cluster statistics are reported in **Supplementary Table 1**).

**Figure 3.**
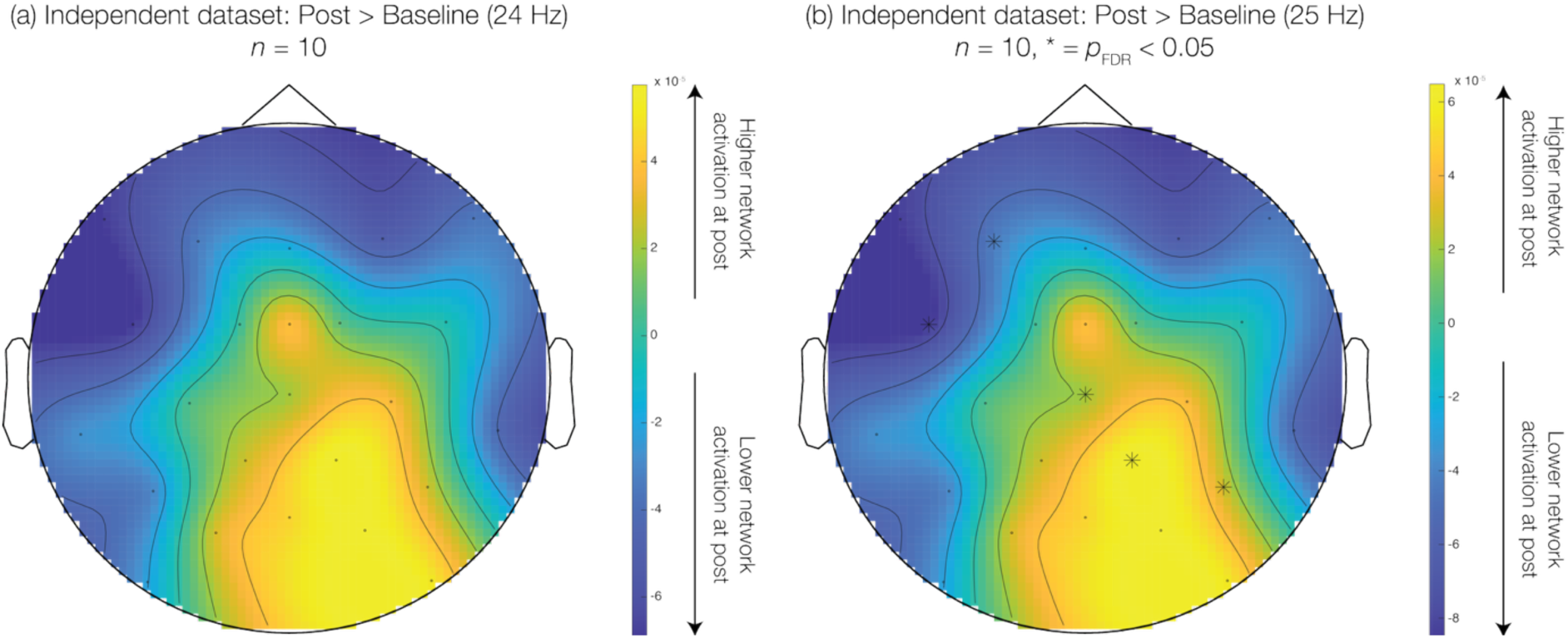
Posterior shifts of high-beta networks after ibogaine were replicated in an independent EEG dataset. Resting-state EEG data were obtained at baseline and after ibogaine from 10 separate participants with opioid use disorder. Ibogaine treatment was associated with a clear posterior shift in the 24 and 25 Hz networks. In the 25 Hz network, decreases in network activation in left frontal electrodes and increases in network activation in posterior electrodes were significant.

### Section 3.3. Posterior shifts in high-beta networks are significantly correlated with PTSD improvements

Next, we investigated the relationship between the posterior high-beta network shift and changes in PTSD symptoms. The position of the electrode along the anterior-posterior axis (the *y*-coordinate) of the brain predicted the correlation of the network activation at that electrode and changes in PTSD symptoms, as measured by the total CAPS-5 score (**Figure 4**). In other words, the more posterior an electrode was, the more positively the electrode’s network activation was associated with PTSD improvement. Likewise, the more anterior an electrode was, the more negatively the electrode’s network activation was associated with PTSD improvement. The association between PTSD improvement and network activation at a particular electrode was significantly correlated with the *y*-coordinate of the electrode at both 24 and 25 Hz (especially the former) and at both timepoint comparisons (24 Hz, immediate-post > baseline: Pearson’s *r* = - 0.66, *p*_corrected_ < 10^-8^; 24 Hz, one month-post > baseline: Pearson’s *r* = −0.53, *p*_corrected_ < 10^-4^; 25 Hz, immediate-post > baseline: Pearson’s *r* = −0.41, *p*_corrected_ = 0.001; 25 Hz, one month-post > baseline: Pearson’s *r* = −0.30, *p*_corrected_ = 0.02). Overall, these results indicate that posterior increases and anterior decreases in network activation after ibogaine are associated with improvements in PTSD symptoms.

**Figure 4.**
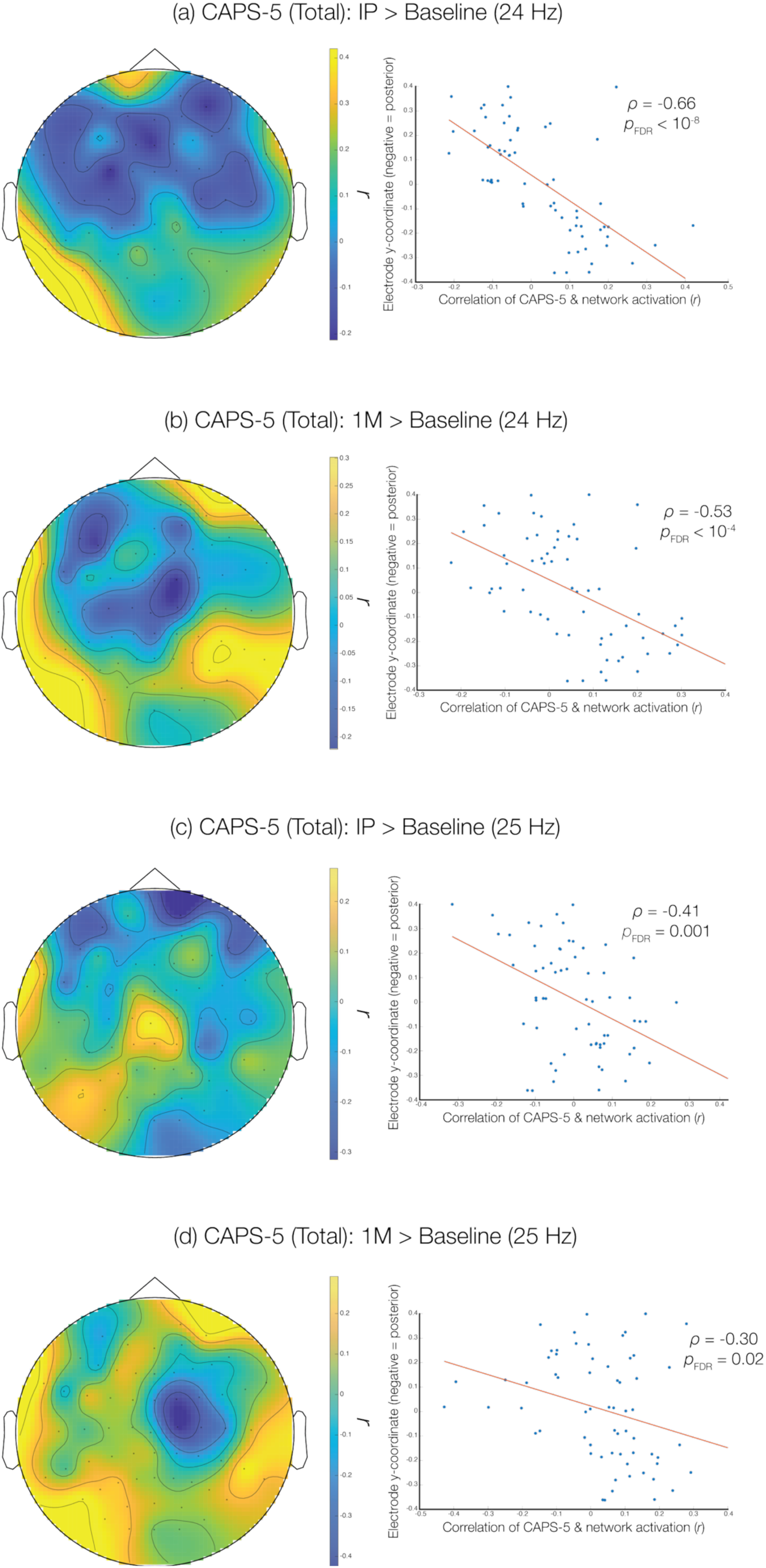
Posterior shifts in the high-beta network topography are significantly correlated with improvements in PTSD symptoms, both immediately and one month after ibogaine. Each electrode’s network activation (the values shown in Figure 2) was correlated with changes in PTSD symptoms, as measured by the total CAPS-5 score. Changes in CAPS scores were multiplied by −1 such that increases in CAPS scores reflected improvement. With Pearson correlations, we then determined whether the strength of this association could be predicted by the *y*-coordinate of the electrode, i.e., its position along the anterior-posterior axis of the brain. *p*-values of each correlation were corrected with FDR across 20 comparisons (2 networks x 2 timepoints x 5 scores, including total CAPS-5 score and four subscale scores [**Supplementary** Figures 5-8]). More posterior electrodes exhibited significantly more positive correlations between their network activation and PTSD improvements, whereas more anterior electrodes exhibited significantly more negative correlations between their network activation and PTSD improvements. IP = immediate-post; 1M = 1 month-post.

We further investigated the association between the posterior high-beta network shift and improvements on specific PTSD subscales. The posterior position of the electrode also predicted the correlation between network activation and scores on specific subscales of the CAPS-5 (**Supplementary Figure 5-8**). This pattern was most evident for subscale B, which measures the intrusiveness of traumatic memories and distress from trauma-related triggers. For subscales D and E, which capture negative alterations in cognition and mood and symptoms of arousal and reactivity, respectively, anterior electrodes tended to exhibit higher correlations between CAPS-5 scores and activation in some of the high-beta networks. In other words, the posterior shift may selectively mediate certain PTSD symptoms. The relationship between the posterior shift and CAPS-5 subscales also appeared to depend on the specific high-beta frequency, as discussed further in **Supplementary Results 4**. Sensitivity analyses demonstrated that this relationship is robust to a different method of measuring the posterior shift (**Supplementary Figure 9**).

### Section 3.4. Model-based cortico-cortical connectivity decreases after ibogaine

We next applied a thalamocortical neural field model (Robinson-Rennie-Wright [RRW]) to determine whether posterior shifts in network topographies arise from changes in thalamocortical connectivity (**Figure 5a**). The RRW model represents large-scale brain dynamics as the interaction of cortical and thalamic neural populations connected by excitatory and inhibitory loops. By fitting the model to EEG power spectra at each timepoint, we can estimate how the effective coupling between these populations changes after ibogaine and then determine which of these changes are sufficient to reproduce the empirical shifts in high-beta network topography. **Figure 5b** displays the fit of the RRW model to the power spectrum of a single electrode of a representative participant. High-beta power (>20 Hz) in the fitted spectrum closely resembled that of the empirical spectrum. (Median goodness-of-fit, or χ^2^, across electrodes and participants at baseline, immediate-post, and one month-post is 1.35, 1.22, and 1.24, respectively.)

**Figure 5.**
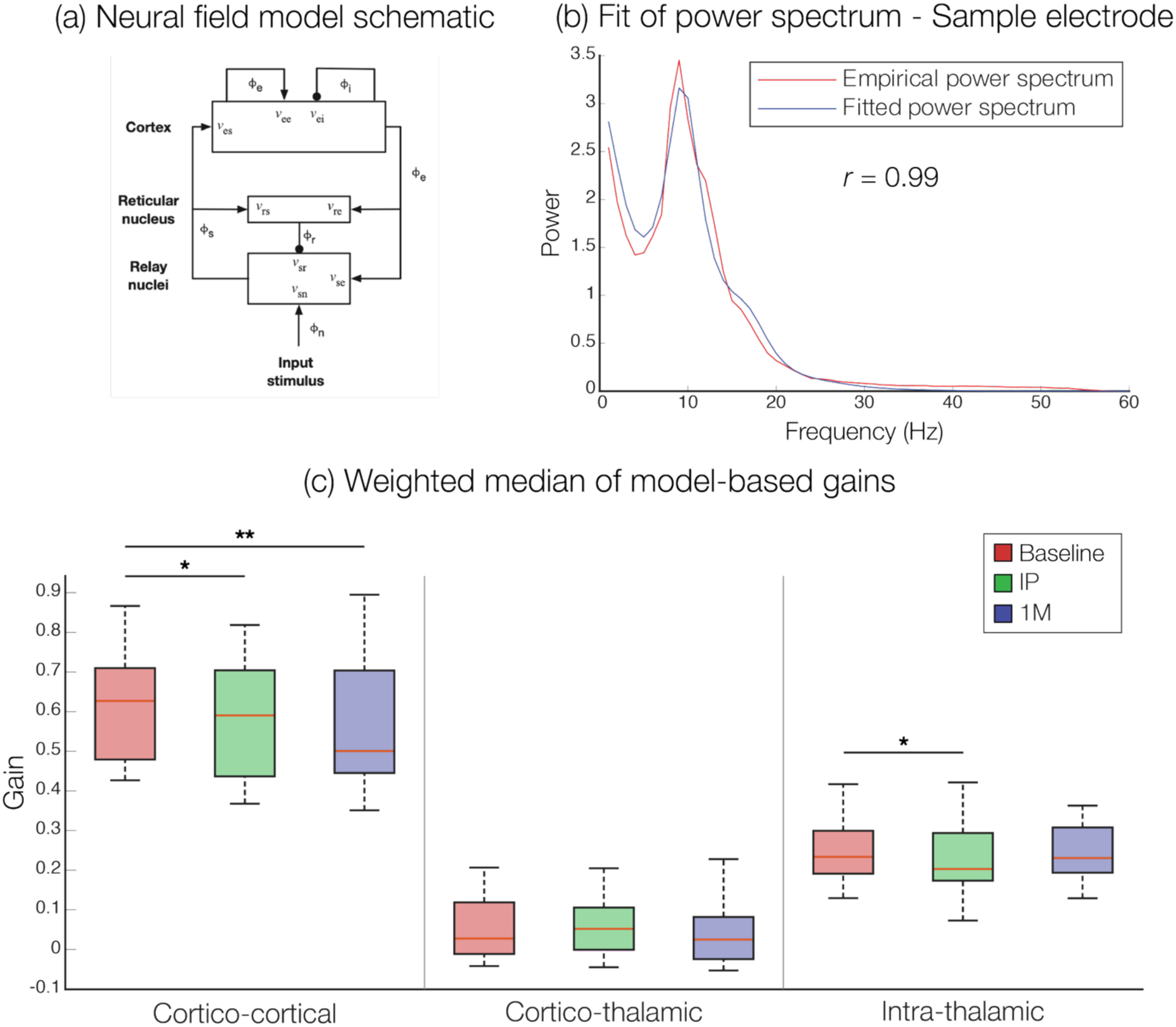
Ibogaine is associated with significant decreases in cortico-cortical connectivity at both immediate-post and 1 month-post timepoints. (a) We used a Robinson-Rennie-Wright corticothalamic neural field model comprising four interacting neural populations: cortical excitatory and inhibitory neurons, thalamic relay neurons, and thalamic reticular neurons. Image is reproduced with permission from Abeysuriya et al. (2014). (b) The steady-state linearized form of the model provided an analytic expression for the EEG power spectrum, which was fitted to empirical spectra. Fitting was performed via BrainTrak, which uses a Metropolis–Hastings Markov Chain Monte Carlo algorithm that minimized the weighted fractional difference between model and experimental power spectra. This plot shows the empirical and fitted power spectra for a single electrode (PO7) from an arbitrary participant; the two spectra are evidently similar. (c) The central dynamics of the model are captured by three parameters representing the cortico-cortical, cortico-thalamic, and intrathalamic gains, respectively. For each subject, the median of each parameter across electrodes was weighted by the inverse goodness of fit (χ^2^). Cortico-cortical gain significantly decreased at both the immediate-post and 1 month-post timepoints. However, ibogaine did not have a significant effect on cortico-thalamic gain at either timepoint. Intrathalamic gain significantly decreased at the immediate-post timepoint but not the 1 month-post timepoint. IP = immediate-post; 1M = 1 month-post.

We measured the effect of ibogaine on model-based cortico-thalamic, cortico-cortical, and intrathalamic gain, which together constitute the core dynamics of the model. Our hypothesis was that ibogaine would specifically and significantly increase the model-based cortico-thalamic gain. However, ibogaine was instead associated with significant decreases in the median of the cortico-cortical gain across electrodes, weighted by χ^2^, at both the immediate-post (β = −0.0376, standard error [SE] = 0.0180, t(53.05) = −2.091, *p* = 0.04) and one month-post timepoints (β = −0.0581, SE = 0.0187, *t*(53.41) = −3.1, *p* = 0.003) (**Figure 5c**). This corresponds to a reduction in effective excitatory coupling across cortex. Contrary to our hypothesis, ibogaine did not have a significant effect on the weighted median of the cortico-thalamic gain at either timepoint (immediate-post: β = 0.0123, SE = 0.0102, *t*(52.97) = 1.205, *p* = 0.23; one month-post: β = 0.0064, SE = 0.0106, *t*(53.36) = 0.605, *p* = 0.55) . The weighted median of the intrathalamic gain was significantly reduced at the immediate-post timepoint (β = −0.0201, SE = 0.0077, *t*(52.93) = −2.625, *p* = 0.01), but not at the one month-post timepoint (β = −0.0101, SE = 0.0080, *t*(53.14) = −1.268, *p* = 0.21).

### Section 3.5. Decreases in cortico-cortical gain are associated with posterior shifts in simulated high-beta networks

Finally, we used neural field simulations to test whether the fitted gain parameters reproduce the empirical high-beta network topographies. Indeed, the empirical 24 Hz network topography at baseline, averaged across subjects, strongly resembled the simulated topography (*r* = 0.82, *p* < 10^-15^), which used the gain parameters fitted to the baseline EEG power spectra (**Figure 6a**, **6c**). Both the empirical and simulated topographies exhibit a clear peak of activation in a left frontal region. In both cases, activation was much lower in posterior electrodes. This correspondence between the simulated and empirical baseline topographies validates the model’s ability to capture the spatial distribution of high-beta network activity, and thereby justifies the use of the model to test which parameter changes reproduce the observed post-ibogaine shifts in network topography. We found that simulated decreases in the cortico-cortical gain yielded network topographies that were similar to the empirical post-ibogaine topographies. Empirically, as noted in *Section 3.1*, ibogaine reduced the activation of frontal left electrodes in the 24 Hz network at the one month-post timepoint (**Figure 6b**). In simulations, decreasing the cortico-cortical gain also led to reductions in left frontal activation in the 24 Hz network (**Figure 6d-f**). We reduced the gain of the connectivity between excitatory cortical and excitatory cortical neurons by intervals of 10% and showed that larger reductions were associated with lower activation in left frontal electrodes. Indeed, the simulated difference between the activation of left frontal and posterior electrodes monotonically grew as the cortico-cortical gain decreased, approximating the empirical difference when the cortico-cortical gain was reduced by just 10-20% (**Figure 6g**). This is consistent with the model-based reduction of 15.46% in cortico-cortical gain between the empirical baseline and 1 month-post data.

**Figure 6.**
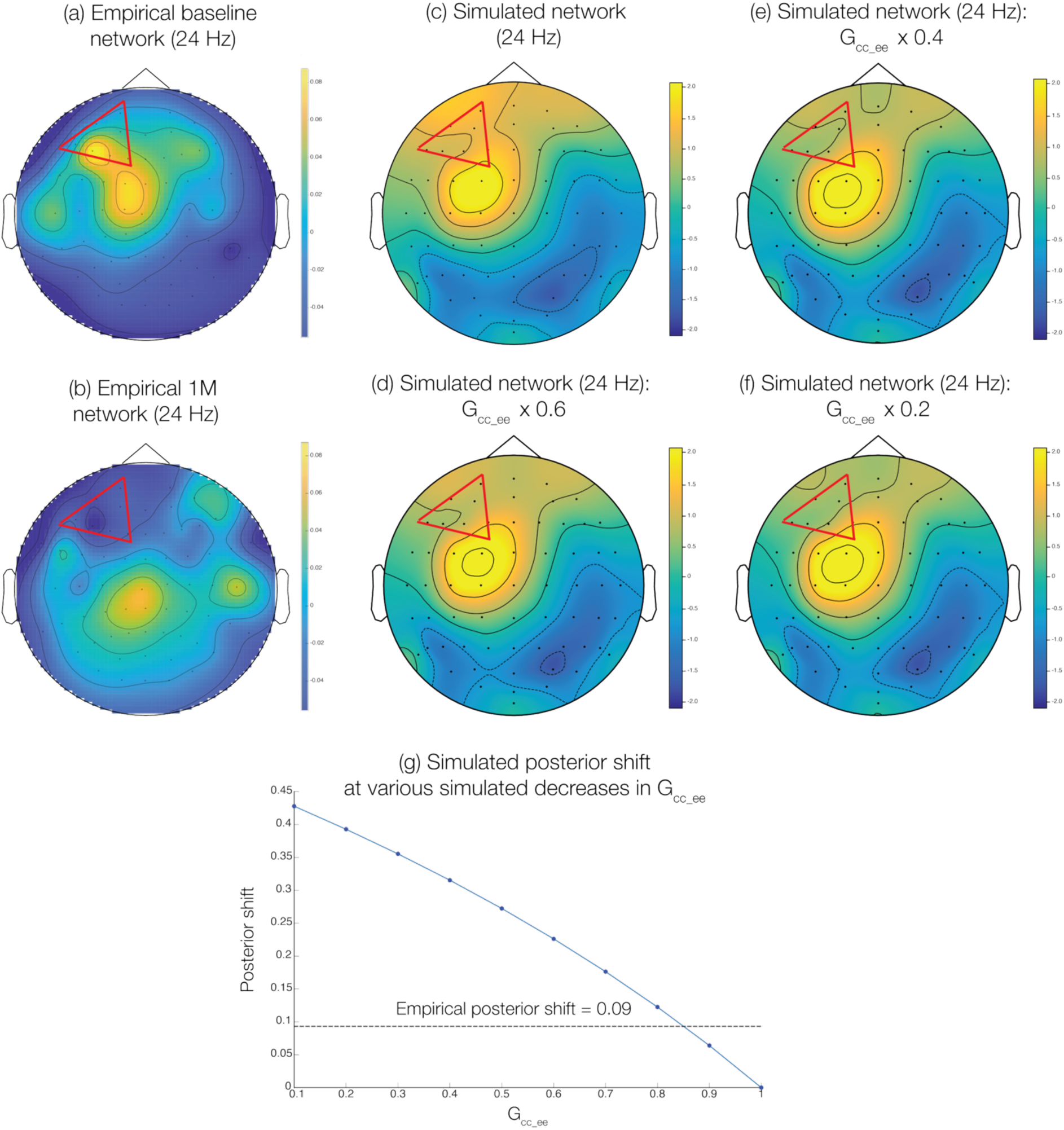
Simulated high-beta network topographies match empirical topographies both before and one month after ibogaine. (a) The empirical group-averaged 24 Hz network topography at baseline shows a clear left frontal peak, as highlighted by the red triangle. (b) The left frontal peak disappears at 1 month-post; ibogaine is associated with substantial reductions in the activation of left frontal electrodes in the 24 Hz network. (c) RRW model parameters that had been fitted with BrainTrak were used to simulate the 24 Hz network topography. Like the empirical network, the simulated network exhibits both a central and left frontal peak. (d-f) Progressively decreasing the cortico-cortical gain (specifically, the excitatory-excitatory gain, i.e., G_cc_ee_) by a factor of 40, 60, and 80%, respectively, is associated with a gradual reduction of frontal left activation in the simulated 24 Hz network. Therefore, decreases in cortico-cortical gain are linked to the loss of the frontal left peak 1 month after ibogaine, in the empirical 24 Hz network. (g) We measured the posterior shift based on the difference in network activation patterns between the left frontal electrodes (those in the red triangle) and posterior electrodes. The empirical posterior shift is denoted by the dashed line. The simulated posterior shift was calculated for a range of G_cc_ee_ values. Reducing G_cc_ee_ by just 10-20% (i.e., multiplying G_cc_ee_ by 0.8-0.9) was sufficient to approximate the empirical posterior shift.

To rule out the possibility that increases in cortico-cortical gain could equally account for the empirical topographic changes, we also simulated the effects of elevating this gain parameter. Although increasing this gain by 40% or higher also caused a loss of left frontal activation (**Supplementary Figure 10**), such increases predicted spectral and connectivity features that were inconsistent with the empirical data. In particular, increases of 30% or more dramatically elevated the simulated beta power, averaged across nodes, whereas ibogaine did not significantly affect mean empirical beta power at either timepoint (**Supplementary Figure 11a-b**). Additionally, increases of 40% or higher in the cortico-cortical gain are associated with a posterior shift that is much greater in magnitude than the empirical posterior shift (**Supplementary Figure 11c**). Therefore, decreases in cortico-cortical gain are consistent with the lack of change in empirical beta power and increases in empirical variance explained by the 24 Hz network.

While ibogaine does significantly decrease the intrathalamic gain at the immediate-post timepoint, reducing this gain in the simulations by as much as 90% did not alter the 24 Hz network topographies (**Supplementary Figure 12**). Thus, changes in intrathalamic gain do not appear to be associated with posterior shifts in high-beta network topographies.

## Section 4. Discussion

This is the first study to report the effects of ibogaine on the organization of brain networks oscillating at specific frequencies, as well as their relationship to clinical outcomes. We found that U.S. veterans with a history of TBI and varying levels of PTSD exhibited significant changes in the topographies of high-beta networks after ibogaine treatment. In particular, activation of these networks decreased in left frontal electrodes and increased in posterior electrodes. This posterior shift was present both 3-4 days and one month after treatment and, critically, correlated with improvements in PTSD symptoms. The posterior shift also replicated in an independent EEG dataset from patients with OUD who were treated with ibogaine. With neural field modeling, we demonstrated that decreases in cortico-cortical connectivity, not cortico-thalamic or intrathalamic connectivity, are associated with some of the observed shifts in high-beta networks.

### Section 4.1. Beta-band networks regulate the symptoms of PTSD and are reorganized after ibogaine

PTSD and trauma are associated with increases in frontal beta or high-beta power, both during the resting state and when participants are listening to a script describing their traumatic event (Begić et al., 2001; Jang et al., 2017; Reuveni et al., 2022). Early life stress, which is a key predictor of adult PTSD, is also linked to increases in the rate of frontal beta activity (Kavanaugh et al., 2024). Meanwhile, decreases in the duration of frontal beta activity or in beta power correlate with improvements in PTSD after treatments such as repetitive transcranial magnetic stimulation and neurofeedback (Kolk et al., 2016; Morris et al., 2023; Rogel et al., 2020).

Rather than facilitating flexible updating, beta activity, particularly in frontal cortex, is thought to reinforce or maintain the current cognitive or emotional state by stabilizing top-down control signals (Antzoulatos & Miller, 2016; Engel & Fries, 2010; Lundqvist et al., 2024; Miller et al., 2018). For instance, prefrontal beta bursts play a role in maintaining specific items in working memory by inhibiting interference from irrelevant stimuli (Bastos et al., 2012, 2020). Compared to healthy controls, patients with PTSD exhibit less suppression of beta power in left orbitofrontal cortex, as well as in other brain regions, when they are encoding new images into their working memory (Popescu et al., 2020). This failure of active suppression may allow trauma-related neural representations, normally kept in check by prefrontal beta-mediated inhibition, to intrude involuntarily into awareness. This intrusion of traumatic memories into the present moment is a hallmark symptom of PTSD (Krans et al., 2009; Meyer et al., 2013, 2017). Indeed, left frontal beta power uniquely predicts symptoms of intrusive trauma-related memories, including flashbacks, in active combatants (Moran et al., 2017). Other studies have also shown that beta activity plays a role in the lack of cognitive flexibility in PTSD (Popescu et al., 2023); this may further mediate symptoms of intrusion, which are typically characterized by inflexible, recurrent representations of trauma.

Thus, the core pathology of PTSD may lie in a beta-mediated “stuck” top-down control system that cannot allow new learning or flexibly release traumatic content, effectively locking the brain into trauma-related representational states. Within this context, the frontal weighting of high-beta networks, as observed at baseline in our cohort, may reflect a pathological over-engagement of frontal control systems. Using FREQ-NESS, we demonstrated that ibogaine significantly redistributes the cortical regions that dominate beta-band covariance, shifting control away from frontal hubs toward posterior cortex. This recruitment of posterior, particularly occipital, regions may indicate a transition towards a regime in which the brain is predicting sensory inputs, rather than rigidly enforcing threat-biased priors generated by frontal control systems. Indeed, occipital beta activity inhibits sensory processing after stimuli disappear, preventing further sensory updating while stimuli get encoded into working memory (Liljefors et al., 2024). Occipital beta also contains top-down signals that aid in higher-order interpretation and contextualization of visual inputs (Michalareas et al., 2016; Richter et al., 2018; Turner et al., 2023).

In other words, the occipital reorganization of high-beta networks after ibogaine may indicate that sensory areas are contributing more to beta-band connectivity than frontal areas involved in the cognitive and affective representation of traumatic memories, enabling sensory evidence to more effectively update maladaptive trauma-related priors. This potential increased sensitivity to sensory information may play a role in impeding traumatic memories from intruding into present awareness. Consistent with the role of frontal beta in maintaining intrusive trauma representations, the posterior shift in high-beta networks correlated most strongly with improvements on the

Intrusion (Criterion B) subscale of the CAPS-5, compared to changes in other PTSD criteria. Other subscales, such as Cognition/Mood (Criterion D) and Arousal/Reactivity (Criterion E), were sometimes anticorrelated with the posterior shifts, depending on the frequency and timepoint, demonstrating that the observed changes in network topography were most consistently associated with Intrusion.

A popular theory, RElaxed Beliefs Under pSychedelics (REBUS), would support ibogaine’s role of updating inflexible trauma-related neural representations, such as intrusive memories (Carhart-Harris & Friston, 2019). REBUS claims that psychedelics decrease the precision-weighting of “priors” encoding rigid, pathological beliefs in psychiatric conditions (Carhart-Harris & Friston, 2019). (Note that REBUS mostly focuses on “classic” psychedelics like LSD and psilocybin.) In PTSD, overly precise priors in frontal areas, i.e., priors that are assigned an exceedingly high degree of confidence, may maintain the brain in a cognitive and emotional state of perpetual threat, preventing the brain from incorporating new evidence that the environment is safe (Kube et al., 2020; Putica et al., 2022). According to REBUS, psychedelics would redistribute oscillatory control away from dysfunctional frontal circuits, such that patients can disengage from trauma-related attractor states by amplifying their sensory awareness of the environment (Girn et al., 2022; Shinozuka, Tewarie, et al., 2025; Singleton et al., 2022). This posterior redistribution of high-beta brain networks was observed not only in our sample of PTSD patients but also in an independent cohort of OUD patients who received ibogaine treatment. Thus, this shift is not specific to individuals with TBI and may reflect a transdiagnostic neural signature of ibogaine treatment (Nicolas, 2005).

It is important to note that FREQ-NESS does not measure beta power but rather the distribution of the regions contributing most strongly to the covariance structure at specific frequencies. Previous analyses showed that ibogaine does not have a significant effect on beta power at the one month-post timepoint (Lissemore, Chaiken, et al., 2025). As we showed here, standard pairwise connectivity measures like wPLI also may not differentiate brain coupling before and after ibogaine. FREQ-NESS is complementary to conventional spectral methods; whereas the latter indicate nonspecific changes in amplitude or pairwise connectivity, FREQ-NESS captures the reorganization of whole-brain, frequency-specific networks.

Overall, these findings indicate that, across multiple psychiatric conditions, ibogaine does not simply suppress pathological beta activity but selectively reshapes the network-level deployment of beta-band control, weakening rigid frontal dominance and enabling posterior regions to contribute more strongly to the flexible updating of internal models.

### Section 4.2. Intracortical connectivity mediates the effects of ibogaine to a greater extent than cortico-thalamic connectivity

A longstanding hypothesis in psychedelic neuroscience is that these compounds exert their characteristic perceptual and emotional effects by altering thalamocortical gating (Avram et al., 2021; Müller et al., 2017; Vollenweider & Geyer, 2001). Under this framework, the thalamus acts as a central hub that regulates the precision and routing of ascending sensory inputs, filtering out irrelevant stimuli (Aguilar & Castro-Alamancos, 2005; McCormick & Bal, 1994; Zare et al., 2023). Psychedelics are thought to “open the doors of perception,” increasing cortical exposure to raw sensory signals that would otherwise be filtered out by thalamic inhibitory circuits (Coleman et al., 2025; Vollenweider & Geyer, 2001). Furthermore, prefrontal beta bursts are thought to arise from loops between the thalamus and other subcortical areas (Lundqvist et al., 2024; Sherman et al., 2016; Wessel & Anderson, 2024). Meanwhile, PTSD is associated with reduced functional connectivity between the thalamus and occipital, sensorimotor, and salience regions (Kearney et al., 2025; Steele et al., 2026). TBI can lead to neuronal loss in several thalamic nuclei, while also disrupting cortical dynamics by injuring thalamocortical connections (Mofakham et al., 2022; Woodrow et al., 2024).

Recent applications of neural field models to resting-state magnetoencephalography (MEG) data also demonstrated that cortico-thalamic connectivity exhibits a clear anterior-posterior gradient, with posterior regions exhibiting higher connectivity compared to anterior areas (Bastiaens et al., 2025). Cortico-cortical connectivity displayed the opposite pattern. Due to the posterior shift that we observed in our data, as well as the prior literature about the effects of psychedelics on thalamocortical connectivity, we hypothesized that ibogaine should increase cortico-thalamic and potentially intrathalamic connectivity. However, our neural field model indicated that ibogaine did not significantly alter cortico-thalamic gain and that even substantial simulated reductions in intrathalamic gain did not reproduce the empirical reorganization of beta-network topographies. Instead, the posterior shifts in high-beta networks were associated exclusively with decreases in cortico-cortical gain, suggesting that ibogaine’s mechanism of action may not primarily involve modulating thalamic gating, at least in the high-beta range.

Importantly, we cannot conclude that thalamic mechanisms are uninvolved in ibogaine’s effects. Rather, our findings indicate that, when specifically fitting the RRW model, we did not observe clear model-based evidence for cortico-thalamic involvement in the high-beta network shifts. Nevertheless, ibogaine, with its complex, multi-receptor profile and prolonged kinetics, may engage thalamic circuits differently from classic psychedelics such as psilocybin or LSD, which are selective for the 5-HT_2A_ receptor (Cherian, Shinozuka, et al., 2024; Glick et al., 1997; Knuijver et al., 2024; Maciulaitis et al., 2008; Shinozuka, Jerotic, et al., 2024). For example, ibogaine is known to be an agonist at the kappa-opioid receptor, which is expressed less densely in the thalamus than the 5-HT_2A_ receptor (**Supplementary Results 6**). Furthermore, ibogaine’s hallmark phenomenology is intensely introspective, narrative-driven, and less perceptually distortive than classic psychedelics (Cherian, Shinozuka, et al., 2024; B. Cohen & Shenk, 2025; Heink et al., 2017; Mash, 2023). Compared to the subjective experience of ibogaine, sensory hallucinations on classic psychedelics may therefore depend more on reductions in thalamic sensory gating. Previous studies showing that classic psychedelics increase cortico-thalamic connectivity recorded brain activity during the acute effects (i.e., during the “trip”), whereas we recorded EEG data before and after, but not during, the acute effects of ibogaine. It is possible that the thalamus may be more involved in hallucinations during the acute effects of psychedelics than in the post-acute recovery and integration (Onofrj et al., 2023).

It is worth noting that the RRW was originally designed to model alpha oscillations, in part because the thalamus plays a central role in generating alpha (Halgren et al., 2019; Hughes & Crunelli, 2005; Schreckenberger et al., 2004). However, RRW can still vary the parameters that broadly shape cortical gain, recurrent excitation, and corticocortical coupling, and these parameters influence all frequency bands, not only alpha (Robinson et al., 2003). The model is therefore appropriate for testing whether changes in global cortical recurrence, i.e., cortico-cortical gain, can reorganize network topographies, even if it does not fully capture the mechanistic biophysics of high-beta generation.

Overall, significant changes in high-beta network topographies after ibogaine are consistent with our finding that ibogaine affected the modeled cortico-cortical gain but not cortico-thalamic gain. Beta-band activity has been repeatedly linked to intracortical communication, whereas alpha rhythms are more tightly coupled to cortico-thalamic resonance (Halgren et al., 2019; Hall et al., 2011; Hughes & Crunelli, 2005; Jensen et al., 2005; Muthukumaraswamy et al., 2013; Rossiter et al., 2014; Schreckenberger et al., 2004). Within this framework, modulation of cortico-cortical gain would be expected to disproportionately impact beta-band network organization, amplifying or reweighting long-range cortical interactions without necessarily engaging thalamic mechanisms. The preferential reconfiguration of high-beta networks under ibogaine may therefore reflect a shift in cortical processes, potentially destabilizing rigid, over-constrained cortical network states, while preserving the integrity of cortico-thalamic dynamics that support basic sensory processing and rhythmic coordination.

### Section 4.3. Limitations and future directions

Some limitations of this study should be acknowledged. First, the observational design and absence of a control group preclude strong causal inferences about ibogaine’s effects on brain networks or PTSD symptoms. Although we observed robust posterior shifts in high-beta networks and significant correlations with symptom improvements, placebo effects, expectancy, and non-specific influences such as the treatment setting cannot be excluded.

Second, although neural field modeling provided mechanistic insight into how cortical and thalamic parameters may shape high-beta network topographies, the RRW framework imposes important constraints on inferences about the thalamus. RRW is a coarse-grained, mass-signal model that represents thalamic relay and reticular nuclei as spatially distributed but functionally homogeneous populations (Abeysuriya et al., 2014; Abeysuriya & Robinson, 2016; Robinson et al., 1997, 2003). It cannot resolve thalamic subnuclei (e.g., pulvinar versus mediodorsal nucleus), laminar-specific projections, or the heterogeneity of cortico-thalamic pathways that are known to shape oscillatory dynamics. Future source-space analyses may be able to identify these contributions, but subcortical activity is generally difficult to directly measure in EEG data (Michel & Brunet, 2019). The model estimates effective loop gains, not direct synaptic or structural connections, and these gains are fit to EEG power spectra. Thus, cortico-thalamic connectivity is being indirectly inferred from the power spectrum. Our modeling should be interpreted as demonstrating that decreases in cortico-cortical gain are sufficient to reproduce the observed network shifts, not that thalamic mechanisms play zero role in the ibogaine experience.

We encourage future researchers to directly link beta-band activity to the symptoms of PTSD. For instance, while recording EEG or MEG activity, researchers could use task-based paradigms to directly link frontal beta activity to intrusive memories or impairments in attention and impulse control among patients with PTSD. This could validate the functional role of beta bursts in the maintenance of default cognitive states that characterize PTSD symptoms.

One final limitation is that frequencies above 30 Hz were not analyzed. Gamma-band activity is difficult to reliably estimate in scalp EEG because it is highly susceptible to muscle artifacts, microsaccades, and environmental noise, and because the spatial smoothing inherent to volume conduction attenuates fast, spatially localized cortical rhythms (Muthukumaraswamy, 2013). However, gamma oscillations are of particular clinical interest in psychedelic research. For example, increases in gamma power and synchrony have been repeatedly associated with the antidepressant effects of ketamine (Farmer et al., 2020; Medeiros et al., 2023; Purohit et al., 2025). Future studies using MEG, intracranial recordings, or carefully controlled high-density EEG optimized for high-frequency analysis may clarify whether gamma-band network reorganization provides complementary or more proximal biomarkers of psychedelic treatment response.

## Section 5. Conclusion

This study identifies a robust, mechanistically interpretable, long-term neural biomarker of ibogaine treatment in a cohort of veterans with TBI and PTSD. Using frequency-resolved network estimation, we show that ibogaine induces a persistent posterior reorganization of high-beta (24-25 Hz) networks, which strongly correlated with improvements in PTSD symptoms. This network-level shift was replicated in an independent cohort of OUD patients who received ibogaine treatment. Neural field modeling further suggests that this reorganization can be explained by a reduction in cortico-cortical gain, consistent with a relaxation of rigid, maladaptive top-down control states. Together, these findings position the posterior shift of high-beta networks as a candidate biomarker of sustained therapeutic change, linking a specific oscillatory mechanism to long-term clinical outcome. More broadly, our results demonstrate that psychedelic interventions can leave enduring signatures in large-scale brain network organization, offering a principled path toward objective markers of treatment response and toward mechanistic models of how psychedelic-assisted therapies promote recovery in trauma-related disorders.

## Acknowledgments

This work was made possible by the generous donations of Steve and Genevieve Jurvetson, the Effie and Wofford Cain Foundation, Eugene Jhong, the Saisei Foundation, Laura Keller, and by support from and collaboration with Veterans Exploring Treatment Solutions (VETS), Inc., a non-profit organization committed to advancing healthcare options for veterans. The funders played no role in study design, execution, data analysis, or preparation of the manuscript.

Several authors of this manuscript are employed by and receive financial support from the WOMEN Center of Excellence and/or the War Related Illness and Injury Study Center at the Veterans Affairs Palo Alto Healthcare System. The contents do not represent the views of the U.S. Department of Veterans Affairs or the United States Government.

We gratefully acknowledge the late Dr. Nolan Williams, who played a central role in the conceptualization and design of this work. Although Dr. Williams passed away prior to manuscript completion and is therefore not included as an author, his scientific vision and contributions were foundational to this study.

## Author Contributions

K.S. conducted all analyses, wrote the manuscript, and conceptualized the project. M.R. guided several analyses, co-developed the FREQ-NESS method, and edited the manuscript. A.C. and J.I.L. preprocessed the EEG data and edited the manuscript. A.C. also source-reconstructed the EEG data for the structure-function coupling analysis. J.I.L. and K.N.C. were responsible for the EEG study design. R.J. conducted the structure-function coupling analysis. N.D. preprocessed the DWI data and extracted the structural connectomes for the structure-function coupling analysis, while also editing the manuscript. V.S. edited the manuscript. M.B. and A.S. acquired the independent EEG dataset and edited the manuscript. K.N.C. performed screening, psychological and cognitive assessments and supervised related scoring and data entry, and led study execution at Stanford for all participants. M.A. edited the manuscript. D.M. advised on the neural field modeling and edited the manuscript. R.D.A. edited the manuscript. L.B. co-developed the FREQ-NESS method and edited the manuscript. M.M.A and C.J.K. edited the manuscript. C.R. supervised the project and edited the manuscript.

## Data and code availability

- Data: The deidentified human participant data reported in this study cannot be deposited in a public repository because they contain sensitive clinical and neuroimaging data from a vulnerable veteran population and are subject to Institutional Review Board-mandated controlled access. To request access, please contact the Stanford University Institutional Review Board and the corresponding author (Cammie Rolle) with a research proposal and data security plan consistent with Stanford data-sharing requirements.
- Code: All code is available upon request by contacting the lead author (Kenneth Shinozuka).

