## Supplementary Materials for "Ibogaine is associated with reorganization of high-beta brain networks in veterans with post-traumatic stress disorder"

### Supplementary Methods

#### Section 1. Participants

The MISTIC clinical trial was preregistered on ClinicalTrials.gov (identifier: NCT04313712). A total of 34 individuals were screened for participation, of whom 30 met eligibility criteria and completed both pre- and immediate-post (3-4 days after ibogaine) treatment assessments. All participants were male Special Operations Veterans (SOVs) with a documented history of traumatic brain injury (TBI) due to blast exposure, head trauma, or combat. The mean age was 44.9 years (s.d. = 7.5). Participants reported an average of 38.6 (s.d. = 52.4) prior TBIs, ranging in severity from mild ($n$ = 28) to moderate ($n$ = 1) and moderately severe ($n$ = 1), as assessed using the Ohio State University TBI Identification Method. Psychiatric diagnoses at study entry, determined via the Mini International Neuropsychiatric Interview (MINI), included PTSD ($n$ = 23), major depressive disorder ($n$ = 15), anxiety disorder ($n$ = 14), and alcohol use disorder ($n$ = 15). In line with treatment safety protocols, participants discontinued medications with potential for negative interactions with ibogaine, including diuretics, CYP2D6-inhibiting medications, serotoninergic medications (that is, any that may increase risk of serotonin syndrome), calcium channel blockers, β-blockers, benzodiazepines, stimulants, corticosteroids and all psychiatric medications.

#### Section 2. Treatment

All participants were referred by the nonprofit organization Veterans Exploring Treatment Solutions, Inc. (VETS), and independently arranged to receive ibogaine treatment at Ambio Life Sciences in Mexico. The MISTIC protocol involved oral administration of ibogaine (mean total dose = 12.1 ± 1.2 mg per kg). One to two hours prior to ibogaine dosing, participants were pre-treated with intravenous magnesium sulfate (1 g) to mitigate cardiovascular risks, particularly Q-T interval prolongation. Prior to dosing, participants underwent medical evaluation and preparatory sessions with a licensed therapist regarding the ibogaine experience. These sessions included intention setting, tools for setting expectations, and psychological preparation for the ibogaine experience, including anxiety management. Integration support was made available following treatment, in which therapists helped patients to process emotions and insights from the ibogaine experience. No psychotherapy was provided during the dosing session itself. Because the psychoactive effects of ibogaine can persist for 24-72 hours or longer, participants remained under continuous medical monitoring for 72 hours post-administration.

#### Section 3. Clinical measures

PTSD symptoms were assessed using the Clinician-Administered PTSD Scale for DSM-5 (CAPS-5). While functional disability, depression, anxiety, and cognitive performance were also measured in the original study, we only correlated changes in brain activity with CAPS-5 scores for several reasons. Firstly, although functional disability was the primary outcome measure of the study, there were some participants who were missing data on functional disability at the one month-post timepoint, whereas CAPS-5 data was collected for all participants at all timepoints. Secondly, more participants were officially diagnosed with PTSD ($n$ = 23) than they were with major depressive disorder ($n$ = 15) or anxiety disorder ($n$ = 14). Thirdly, we sought to avoid multiple comparisons, especially given that changes in brain networks were measured at many frequencies.

**Section 4. EEG data acquisition**

As reported previously (Lissemore et al., 2025), a total of six minutes of eyes-open EEG were recorded, divided into two three-minute blocks separated by a short rest to help participants stay alert. EEG was collected with eyes open to maintain wakefulness and because eyes-open spectral features have been found to reliably predict psychiatric treatment responses. During recording, participants sat upright, fixated on a central crosshair, remained still, and allowed their thoughts to flow naturally. Signals were sampled at 10 kHz from 64 Ag–AgCl electrodes (actiCAP slim; Brain Products GmbH) connected to an actiCHamp amplifier, with Cz serving as the reference channel. Channels were arranged according to the 10–10 system. Impedance was maintained at <10 kΩ in >80% of channels and <25 kΩ in >99% of channels.

**Section 5. EEG preprocessing**

EEG was preprocessed as previously reported, with the exception that the data were downsampled to 200 Hz, rather than 1 KHz, to reduce the amount of data and improve the numerical stability of the FREQ-NESS computations (Lissemore et al., 2025). EEG preprocessing was carried out offline in MATLAB using a combination of the EEGLAB toolbox and the ARTIST automated artifact rejection algorithm. The pipeline included the following steps: (1) 60 Hz line noise and its harmonics were removed with a notch filter, and 1-50 Hz bandpass filtering was applied using windowed-sinc FIR filters; (2) the start and end of each recording were trimmed to avoid filter edge artifacts; (3) bad channels were identified semi-automatically – based on impedance greater than 25 kΩ (consistent with modern high-input-impedance amplifiers), bridging, or persistent noise detected by ARTIST (Wu et al., 2018) – and confirmed by visual inspection (fewer than 15% of channels were excluded). For four datasets, a higher impedance cutoff of 50 kΩ was used to keep rejection within this limit, and visual inspection confirmed good signal quality. Missing channels were replaced using spherical interpolation. (4) The data were segmented into 2-second epochs, and noisy epochs were discarded (mean ± SD = 6.1 ± 2.6%; range = 1.7-13.9%). Epochs were then concatenated together. (5) Data were re-referenced to the average. (6) Independent component analysis was performed using the Infomax algorithm with dimension reduction via principal components analysis (retaining the minimum number of components explaining > 99.9% of the variance; mean ± SD = 50.2 ± 4.7 components, range = 35-59). Components corresponding to eye and muscle artifacts were identified and removed based on their time courses, spectra, and spatial topographies. There were no significant differences in the number of components removed across study timepoints (baseline: 6.2 ± 3.3 components; immediate-post: 5.5 ± 2.8 components; one month-post: 7.1 ± 4.8 components). (7) Data were again re-referenced to the average since ICA can alter the global mean of the EEG data. All preprocessing was done by an experienced rater who was blinded to session order and clinical information. (8) As mentioned earlier, data were downsampled to 200 Hz.

**Section 6. Statistical tests on FREQ-NESS**

We used linear mixed-effects models (LMEs) to statistically test changes in the leading eigenvalue, with Timepoint, Frequency, and their interaction as within-subject fixed effects. Subject was included as a random intercept to account for repeated measurements within individuals. *p*-values for frequency-specific effects between pairs of timepoints were corrected for multiple comparisons using the false discovery rate (FDR) procedure. LMEs were fit with the lmer function in R.

For cluster-based permutation testing on the network topographies (i.e., activation patterns corresponding to leading eigenvectors) associated with each discrete frequency, two-tailed paired *t*-tests were applied to network activation patterns at each electrode, spatial clustering was performed across neighboring electrodes, and cluster-level significance was evaluated using a Monte Carlo permutation procedure (5,000 randomizations; cluster-forming threshold $\alpha = 0.05$; maximum cluster size statistic). For each frequency and contrast, the minimum cluster-level *p*-value (across positive and negative clusters) was retained. To control for multiple comparisons across frequencies and contrasts (29 frequencies x 2 contrasts), all cluster-level p-values were adjusted with FDR.

**Section 7. Weighted phase lag index**

Weighted phase lag index (wPLI), $\Phi$, was calculated only on the frequencies that were associated with significant differences in network topographies. This was done by first bandpass-filtering the signal to the relevant frequencies and then calculating Φ based on the method developed by Vinck and colleagues (Vinck et al., 2011).

Both the network-based statistic (NBS) (Zalesky et al., 2010) and FDR were used to correct wPLI for multiple comparisons. Analyses were restricted to the 27 participants with complete EEG data across all three timepoints. A one-way repeated-measures ANOVA design was specified, with a 27 × 3 one-hot design matrix coding the three timepoints and an exchange-block vector identifying repeated measurements from the same participants. For NBS, a cluster-forming threshold of $F = 3.11$ ($p < 0.05$, uncorrected) was applied, and 5,000 permutations were used to estimate the empirical null distribution of maximal network extent. Significant connected components were identified at a corrected $\alpha = 0.05$ (family-wise error rate).

**Section 8. Analyses to control for the influence of muscle and ocular artifacts on FREQ-NESS results**

To evaluate whether shifts in network topographies under ibogaine were attributable to muscle and ocular artifacts, we ran three control analyses.

Firstly, for each reference scheme (average, linked-ears [see below]), timepoint, and frequency (24, 25 Hz), we used ordinary least squares linear regression to model the relationship between the change in network topography and a proxy of electromyography (EMG) activity, which was the average 40-80 Hz power over several frontal and temporal electrodes (Fp1, Fp2, AF7, AF8, F7, F8, FT7, FT8, T7, T8, FT9, FT10, TP9, TP10) (Goncharova et al., 2003). We then log-transformed and averaged these channel values to obtain a scalar EMG proxy.

Secondly, an electrooculography (EOG) proxy was computed as the *z*-scored log of the standard deviation of the amplitude of Fp1, Fp2, F7, and F8 in each 2-second epoch of data (Yao et al., 2024). We excluded the epochs in the top $20^{th}$ percentile of EOG proxies, recomputed FREQ-NESS, and re-tested shifts in network topographies. Paired Wilcoxon signed-rank tests were used to assess whether exclusion of these epochs changed the posterior shift of the network topography, as determined by subtracting the network activation patterns in the posterior half of the EEG from those of the anterior half.

Thirdly, we recomputed network topographies by changing the EEG referencing scheme from average to linked-ears (TP9/TP10), since EEG data that are referenced close to the ear are more susceptible to EMG artifacts (Goncharova et al., 2003). Paired Wilcoxon signed-rank tests were used to compare the posterior shift between the average-referenced and linked-ears-referenced EEG data.

#### Section 9. Simulating network topographies

*Section 9.1. Simulation parameters*

NFTsim was used to simulate network topographies based on the parameters fitted by BrainTrak (Sanz-Leon et al., 2018). We modelled the neural field as a 2D square grid consisting of 30 nodes on each side (900 total nodes). The matrix shown in **Figure 4a** was used to model connections between five populations of neurons: the four mentioned above (*e*, *i*, *r*, *s*) and a stimulus population, i.e., a population with no inputs from other neurons. The stimulus population was modelled as white noise with mean of 1 and amplitude spectral density of approximately $7 * 10^{-5}$, in accordance with prior models (Abeysuriya et al., 2014). The remaining neuron populations were modelled with a sigmoidal firing response consisting of the following fixed parameters – maximum firing rate = 340/s, firing threshold = 12.9 mV, threshold spread = 3.8 mV – based on prior literature (Abeysuriya & Robinson, 2016). Initial firing rate, $Q$, varied by population, in line with prior models (Abeysuriya et al., 2014).

Based on prior literature (Abeysuriya et al., 2014), we modelled the propagation of activity from excitatory cortical to excitatory cortical neurons, to inhibitory cortical neurons, to thalamic reticular neurons, and to thalamic relay neurons as waves governed by the equation:

$$\left[\frac{1}{\gamma_{ab}^2}\frac{d^2}{dt^2}+\frac{2}{\gamma_{ab}}\frac{d}{dt}+1-r_{ab}^2\nabla^2\right]Q_b(r,t)=Q_b(r,t-\tau_{ab})$$

where $\gamma = 116/\text{s}$ is the damping rate and $r = 86$ mm is the axon range. For the excitatory cortical to thalamic reticular propagation, and for the excitatory cortical to thalamic relay propagation, $\tau$ is the corticothalamic loop delay, as fit by BrainTrak. Otherwise, $\tau$ is simply set to zero, in accordance with prior literature (Abeysuriya et al., 2014). Propagations of activity between the other populations of neurons were modelled as “maps,” i.e., point-to-point connections with no spatial spread.

Based on parameters fitted with BrainTrak, specific gains were calculated as follows:

$$G_{es}=G_{se}=\sqrt{G_{ese}}$$

$$G_{sr}=G_{rs}=\sqrt{G_{srs}}$$

$$G_{re}=\frac{G_{esre}}{G_{es}*G_{sr}}$$

*Section 9.2. Sensor geometry and forward model*

We used an EEG forward operator $F$ computed on the fsaverage head model (3-layer BEM; ico-5 source space) with BioSemi-64 electrode positions. The forward was converted to fixed, surface-normal orientation so each cortical vertex contributes a single dipolar component.

*Section 9.3. Reference projection on the gain*

To ensure the simulations and empirical data share the same reference, we applied the average-reference projector to the gain,

$$R = I_M - \frac{1}{M}\mathbf{1}\mathbf{1}^{\mathrm{T}}$$

$$F_{\mathrm{AR}} = RF$$

where $M = 64$ is the number of electrodes, so any simulated sensor data $y = Fx$, where $x$ is the simulated 2D grid of the neural field, is implicitly average-referenced as $Ry = F_{\mathrm{AR}}x$.

*Section 9.4. Vertex-centered source packing*

Embedding a two-dimensional simulated neural field onto the three-dimensional brain can introduce an artificial "meridian seam," analogous to folding a flat sheet into a cylinder, which creates a seam from joining the opposite edges of the sheet. To embed the simulated neural field onto cortex without introducing this seam, we:

1. computed spherical angles $(\theta, \phi)$ for each cortical node in head coordinates ($\theta$ is like latitude, in which $\theta = 0$ is aligned with the superior (z-axis) direction in MRI space, and $\phi$ is like longitude)
2. restricted the source space to a superior cap (default < 80°), since EEG is most sensitive to neurons that are oriented radially (perpendicular) to the scalp surface, which tend to be neurons in superior cortex (Goldenholz et al., 2009)
3. selected exactly 900 cortical nodes once (uniform at random within the cap), since there are 900 nodes in the neural field
4. ordered the nodes into 30 bins based on their angular distance from $\theta = 0$
5. within each bin, ordered the nodes by $\phi$, such that adjacent bins are ordered in opposite directions (even-numbered bins increase in $\phi$ and odd-numbered bins decrease in $\phi$)

This last step avoids a global $\phi$ seam by ensuring that the angular wrap-around between $\phi = -\pi$ and $\phi = \pi$ occurs locally within individual bins and does not align across bins to form a global discontinuity in the two-dimensional neural field. After the columns of $F_{\mathrm{AR}}$ are re-ordered by this method and $F_{\mathrm{AR}}$ is multiplied by $x$, high-beta network topographies were computed via FREQ-NESS.

**Section 10. Acquisition of independent EEG dataset**

Eyes-closed resting-state EEG was recorded using 28 Ag/AgCl scalp electrodes mounted in an elastic cap (ActiCap, Brain Products, Munich, Germany) and connected to a QuickAmp EEG system (Brain Products GmbH, Munich). Signals were referenced to the left mastoid, with the ground electrode positioned on the left side of the nose. Channels were arranged according to the 10–10 system. Data were acquired at a sampling rate of 1000 Hz, with no online filtering applied.

Vertical electro-oculogram (EOG) activity was recorded bipolarly using electrodes placed 3 mm above the left eyebrow and 1.5 cm below the lower eyelid, while horizontal EOG was recorded from electrodes positioned 1.5 cm lateral to the outer canthus of each eye. Electrode impedances were maintained below 5 kΩ at the beginning of the session and monitored throughout data collection.

Unlike in the MISTIC EEG dataset, certain frontal electrodes (e.g., AF, Fp) were not recorded. Additionally, recordings from occipital electrodes were flat in multiple participants and therefore excluded from all analyses.

**Section 11. Structure-function (S-F) coupling**

At baseline, diffusion-weighted imaging (DWI) data was acquired from 23 of the participants in our sample. These were used to construct structural connectomes for the S-F coupling analysis. Preprocessing was performed using *QSIPrep* 1.0.2.dev0+gfc89945.d20250405 (Cieslak et al., 2021). A T1w-reference map was computed after registration of the available T1w images using antsRegistration [ANTs 2.4.3]. The anatomical reference image was reoriented into AC-PC alignment via a 6-degree of freedom transform extracted from a full affine registration to the MNI152NLin2009cAsym template. A full nonlinear registration to the template from AC-PC space was estimated via symmetric nonlinear registration (SyN) using antsRegistration. Brain extraction was performed on the T1w image using SynthStrip (Hoopes et al., 2022) and automated segmentation was performed using SynthSeg (Billot et al., 2023) from FreeSurfer version 7.3.1. For diffusion data preprocessing, any images with a b-value less than 100 s/mm^2 were treated as a b=0 image. MP-PCA denoising as implemented in MRtrix3's dwidenoise (Veraart et al., 2016) was applied with an auto-voxel window. When phase data were available, this was done on complex-valued data. After MP-PCA, Gibbs unringing was performed using MRtrix3's mrdegibbs (Kellner et al., 2016). Following unringing, the mean intensity of the DWI series was adjusted so all the mean intensity of the b=0 images matched across each separate DWI scanning sequence. B1 field inhomogeneity was corrected using dwibiascorrect from MRtrix3 with the N4 algorithm (Tustison et al., 2010) after corrected images were resampled. Both distortion groups were then merged into a single file, as required for the FSL workflows. FSL's eddy was used for head motion correction and Eddy current correction (Andersson & Sotiropoulos, 2016). Eddy was configured with a q-space smoothing factor of 10, a total of 5 iterations, and 1000 voxels used to estimate hyperparameters. A linear first level model and a linear second level model were used to characterize Eddy current-related spatial distortion. q-space coordinates were forcefully assigned to shells. Field offset was attempted to be separated from subject movement. Shells were aligned post-eddy. Eddy's outlier replacement was run (Andersson et al., 2016). Data were grouped by slice, only including values from slices determined to contain at least 250 intracerebral voxels. Groups deviating by more than 4 standard deviations from the prediction had their data replaced with imputed values. Data was collected with reversed phase-encode blips, resulting in pairs of images with distortions going in opposite directions. FSL's TOPUP (Andersson et al., 2003) was used to estimate a susceptibility-induced off-resonance field based on b=0 images extracted from multiple DWI series with reversed phase encoding directions. The TOPUP-estimated fieldmap was incorporated into the Eddy current and head motion correction interpolation. Final interpolation was performed using the jac method. Several confounding time-series were calculated based on the preprocessed DWI: framewise displacement (FD) using the implementation in Nipype (Power et al., 2014). The head-motion estimates calculated in the correction step were also placed within the corresponding confounds file. Slicewise cross correlation was also calculated. The DWI time-series were resampled to ACPC, generating a preprocessed DWI run in ACPC space with 1.5mm isotropic voxels. Reconstruction was performed using *QSIRecon* 1.1.2.dev0+gfa8673957.d20250819 (Cieslak et al., 2021). The 156-parcel Schaefer Supplemented with Subcortical Structures (4S) atlas (Glasser et al., 2013; King et al., 2019; Najdenovska et al., 2018; Pauli et al., 2018; Schaefer et al., 2018) was used in the workflow. Cortical parcellations were mapped from template space to DWIS using the T1w-based spatial normalization. Multi-tissue fiber response functions were estimated using the dhollander algorithm. Fiber orientation distributions (FODs) were estimated via constrained spherical deconvolution (Tournier et al., 2004, 2008) using an unsupervised multi-tissue method (Dhollander et al., 2016, 2019). Reconstruction was done using MRtrix3 (Tournier et al., 2019). FODs were intensity-normalized using mtnormalize (Raffelt et al., 2017). Individual, rather than group-averaged, connectomes were used for the S-F coupling analysis. We quantify the structural relationship between pairs of ROIs as the density of the fibers, i.e., the sum of the fibers connecting two regions divided by the region volumes.

S-F coupling measures the alignment between functional activity and the cortico-cortical anatomical pathways. For a physiologically grounded characterization, cortical sources of the EEG are estimated on the same parcellation scheme as the structural connectome. Structural MRIs were acquired with a 3 Tesla GE Discovery MR750 scanner with a 32-channel head-neck imaging coil. GE's BRAVO sequence was used (3D, T1-weighted, FOV=256x256mm; matrix=256x256 voxel; TR=6.39ms, TE=2.62ms, slice thickness=0.9mm, flip angle=12°). MRIs were processed using FreeSurfer (recon-all pipeline) to obtain individual cortical surface reconstructions and tissue segmentations. From these segmentations, a three-layer Boundary Element Model (BEM) was constructed, consisting of the scalp (head), outer skull, and inner skull surfaces. EEG sensor positions were manually coregistered to the individual head surface within Brainstorm to ensure accurate alignment between sensors and anatomy. The forward model was computed using OpenMEEG, assuming a realistic head geometry and a surface-based source space constrained to the cortical mantle (Gramfort et al., 2010; Kybic et al., 2005). A surface source grid was defined on the pial surface with unconstrained dipole orientations (three orthogonal components per vertex). For source reconstruction, we used the Minimum Norm Estimate (MNE) with depth weighting enabled. Noise normalization was performed using a diagonal noise covariance matrix, and regularization was set using a fixed signal-to-noise ratio (SNR = 3; noise regularization = 0.1). The resulting imaging kernel mapped sensor-level EEG activity to current density estimates at each cortical vertex. Regions of interest (ROIs) were registered from the 4S atlas to the subject's surface mesh. To obtain regional time series, unconstrained source estimates within each region of interest (ROI) were flattened using principal component analysis (PCA), and the first principal component was retained as the representative ROI time series. This approach captures the dominant variance across dipole orientations while reducing dimensionality and improving signal robustness.

S-F coupling was measured with previously established methods; the reader is referred to the reference text for the complete mathematical description of the method (Subramani et al., 2025). In brief, harmonics of the structural connectome are obtained by performing eigendecomposition. Lower-order eigenmodes represent patterns of structural connectivity with lower spatial frequencies (i.e., smoother patterns), whereas higher-order eigenmodes represent patterns with higher spatial frequencies (i.e., less smooth patterns). When the functional EEG data is projected onto the connectome using eigenmodes, lower-order eigenmodes reflect functional activity that is constrained to the connectome, whereas higher-order eigenmodes reflect functional activity that is more spatially fragmented and more weakly aligned with the connectome. The structural-decoupling index (SDI) is then the ratio between activity captured by lower-order eigenmodes (coupled components) and activity captured by higher-order eigenmodes (decoupled components) (Preti & Van De Ville, 2019). Here, the regional, functional EEG data was represented as the product between the network activation timeseries $\mathbf{y}$ and the spatial network activation patterns

**a**, after FREQ-NESS was applied to the source-space data. (Note that the results of FREQ-NESS were similar between the source-space and sensor-space data.)

SDI was measured regionally and globally by averaging across regions. To assess statistically significant differences in regional SDI, we performed an LME with Timepoint and Region as fixed effects and Subject as a random effect. Timepoint was the sole fixed effect, with Subject as a random effect, in the LME for global SDI. In both the global and regional cases, FDR was applied to correct for multiple comparisons across frequencies.

**Section 11. Comparison of kappa-opioid receptor (KOR) to 5-HT$_{2A}$ receptor expression**

To compare thalamic expression of the kappa-opioid receptor gene (OPRK1) and the 5-HT$_{2A}$ receptor gene (HTR2A), we queried the Allen Human Brain Atlas (AHBA) via the Allen Brain Map API (api.brain-map.org). Expression data were obtained from the HumanMA human microarray dataset using the `human_microarray_expression` service for the thalamus (Structure ID 4392).

All DNA probes annotated to each gene were included (OPRK1: 3 probes; HTR2A: 25 probes). For each thalamic tissue sample, gene-level expression was computed as the mean across that gene's probes. We then summarized expression across all thalamic samples and across donors. Between-gene comparison was performed on matched samples (paired by sample) using descriptive paired differences and a paired *t*-statistic.

### Supplementary Results

**Supplementary Results 1. Effect of ibogaine on the prominence of frequency-specific brain networks**

We define the "prominence" of frequency-specific brain networks based on their leading eigenvalue, which reflects the relative expression of narrow-band covariance to broadband covariance captured by the corresponding network. After multiple comparisons, our LME revealed no significant effect of ibogaine on the leading eigenvalue associated with any of the 29 frequencies that were tested (**Supplementary Figure 1**). However, the decrease from baseline to immediate-post in the leading eigenvalue associated with the 9 Hz network trended towards significance ($\beta$ = -1.51, SE = 0.64, $t(2582) = -2.37$, $p = 0.0536$). Decreases from immediate-post to one month-post in the leading eigenvalues associated with the 11 and 12 Hz networks also trended towards significance (11 Hz: $\beta$ = -1.44, SE = 0.65, $t(2582) = -2.20$, $p = 0.0827$; 12 Hz: $\beta$ = -1.43, SE = 0.65, $t(2582) = -2.19$, $p = 0.0854$).

**Supplementary Results 2. Sensitivity analysis of changes in frequency-specific network topographies**

One participant exhibited disproportionately high network activation values across frequencies, as observed by visual inspection. When this outlier participant was removed from the analysis, changes in network topographies became significant at three other frequencies – 12, 13, and 23 Hz – while remaining significant at 24 and 25 Hz (**Supplementary Figure 2**). Shifts in the high-alpha (12 and 13 Hz) network topographies were only significant at the immediate-post timepoint, whereas the shift in the 24 Hz network topography was only significant at the 1 month-post timepoint. Changes in both the 12 and 13 Hz network topographies were strongly lateralized, with pronounced increases in activation in the left hemisphere and decreases in the right hemisphere. The high-beta (23 Hz) network topography exhibited a similar pattern to the other high-beta networks, with significant reductions in frontal left and central activation but increases in posterior activation.

**Supplementary Results 3. Weighted phase lag index**

When FDR was used to correct for multiple comparisons across connections, there were no significant changes in wPLI at 25 Hz, but there was a single pair of electrodes, FT10-F8, that exhibited significant differences in wPLI at 24 Hz ($F_{2,24} = 4.88$, $p = 0.02$) (**Supplementary Figure 4a-b**). Post-hoc comparisons showed that ibogaine significantly decreased FT10-F8 wPLI at the immediate-post timepoint ($p = 0.02$), while increasing it at the one month-post timepoint ($p =$

0.0061), relative to baseline. However, changes in FT10-F8 wPLI were not correlated with improvements in PTSD (immediate-post: $\varrho_{Spearman} = 0.25$, $p_{corrected} = 0.42$; one month-post: $\varrho_{Spearman} = 0.21$, $p_{corrected} = 0.59$) (**Supplementary Figure 4c-d**).

**Supplementary Results 4. Association between posterior shift and symptoms on specific CAPS-5 subscales**

Correlations between the posterior shifts of high-beta network topographies and specific CAPS-5 subscales were evaluated statistically with two methods. The first was a simple FDR correction across the *p*-values associated with all 20 comparisons (2 networks x 2 timepoints x 5 CAPS-5 scores, including total score). For the Intrusion (B) and Avoidance (C) subscales, all of the comparisons were significant and negative, except the immediate-post vs. baseline comparison at 24 Hz for subscale C (**Supplementary Figure 5-6**). In other words, the more posterior the electrode (i.e., the more negative the electrode's position along the anterior-posterior axis), the more positively the electrode's network activation was correlated with the subscale B and C scores. However, for the Cognition/Mood (D) and Arousal/Reactivity (E) subscales, certain comparisons were significantly positive, such as the immediate-post vs. baseline comparisons at 25 Hz for subscale D and the one month-post vs. baseline comparisons at 24 and 25 Hz for subscale E (**Supplementary Figure 7-8**). That is, the more *anterior* the electrode (i.e., the more negative the electrode's position along the anterior-posterior axis), the more positively the electrode's network activation was correlated with the subscale D and E scores. The one month-post vs. baseline comparison at 24 Hz for subscale D was still significantly negative.

The second statistical test was an LME to determine whether the association between posterior shift and CAPS-5 scores varied by subscale and frequency. Consistent with the prior statistical approach, the LME revealed a significant main effect of electrode anterior-posterior position ($F_{1,62} = 38.11$, $p < 10^{-7}$), with more anterior electrodes exhibiting less positive correlations between network activation change and CAPS-5 scores. There was a significant main effect of subscale ($F_{3,946} = 15.26$, $p < 10^{-8}$), but no main effect of frequency band ($F_{1,946} = 0.22$, $p = .641$).

The two-way interaction between electrode position and subscale was highly significant ($F_{3,946} = 108.83$, $p < 10^{-15}$), indicating that the anterior-posterior gradient varied substantially across symptom subscales. Neither the electrode position × frequency interaction ($F_{1,946} = 0.20$, $p = .651$) nor the subscale × frequency interaction ($F_{3,946} = 1.44$, $p = .231$) was significant. Critically, the three-way interaction between electrode position, subscale, and frequency was significant. A likelihood ratio test revealed that the three-way model fit the data significantly better than a reduced model with no three-way interaction ($\chi^2(3) = 12.64$, $p = .005$). A Type III ANOVA also indicated that the subscale-dependent anterior-posterior gradient itself varied across frequency bands $F_{3,946}$ = 4.18, $p$ = .006). For example, the anterior-posterior gradient for CAPS-5 Criterion D (Cognition/Mood) scores reversed between the 24 Hz and 25 Hz networks.

**Supplementary Results 5. S-F coupling**

Finally, we investigated whether posterior beta shifts arise from decreased coupling of functional brain activity with the structural connectome, which has been observed in other studies on psychedelics (Luppi et al., 2021, 2023). Ibogaine did not have a significant effect on the Structural-Decoupling Index (SDI), when averaged across regions, in either the 24 or 25 Hz network timeseries (**Supplementary Figure 13a-b**). However, regionally, ibogaine significantly decreased the SDI of the 24 Hz network in the left inferior occipital cortex at the immediate-post timepoint ($p_{FDR}$ = 0.0007), while significantly increasing the SDI of the 24 Hz network in the left superior frontal gyrus at the one month-post timepoint ($p_{FDR}$ = 0.0409) (**Supplementary Figure 13e-f**). No significant regional effects were found for the SDI of the 25 Hz network.

**Supplementary Results 6. KOR vs. 5-HT$_{2A}$ receptor gene expression in the thalamus**

A total of 209 thalamic samples from 6 donors were available for both genes. Mean thalamic expression was higher for HTR2A than for OPRK1 (4.82 vs 3.79, respectively), corresponding to an HTR2A/OPRK1 mean ratio of 1.269. The mean paired difference (HTR2A - OPRK1) across samples was 1.0222, and 78.9% of samples showed higher HTR2A than OPRK1 expression. The paired t-statistic for the sample-wise difference was 11.786.

At the donor level, all six donors showed higher thalamic HTR2A than OPRK1 expression (donor-level deltas range: +0.7689 to +1.7228). Together, these analyses indicate that in AHBA thalamic tissue, HTR2A mRNA expression is consistently greater than OPRK1 mRNA expression.

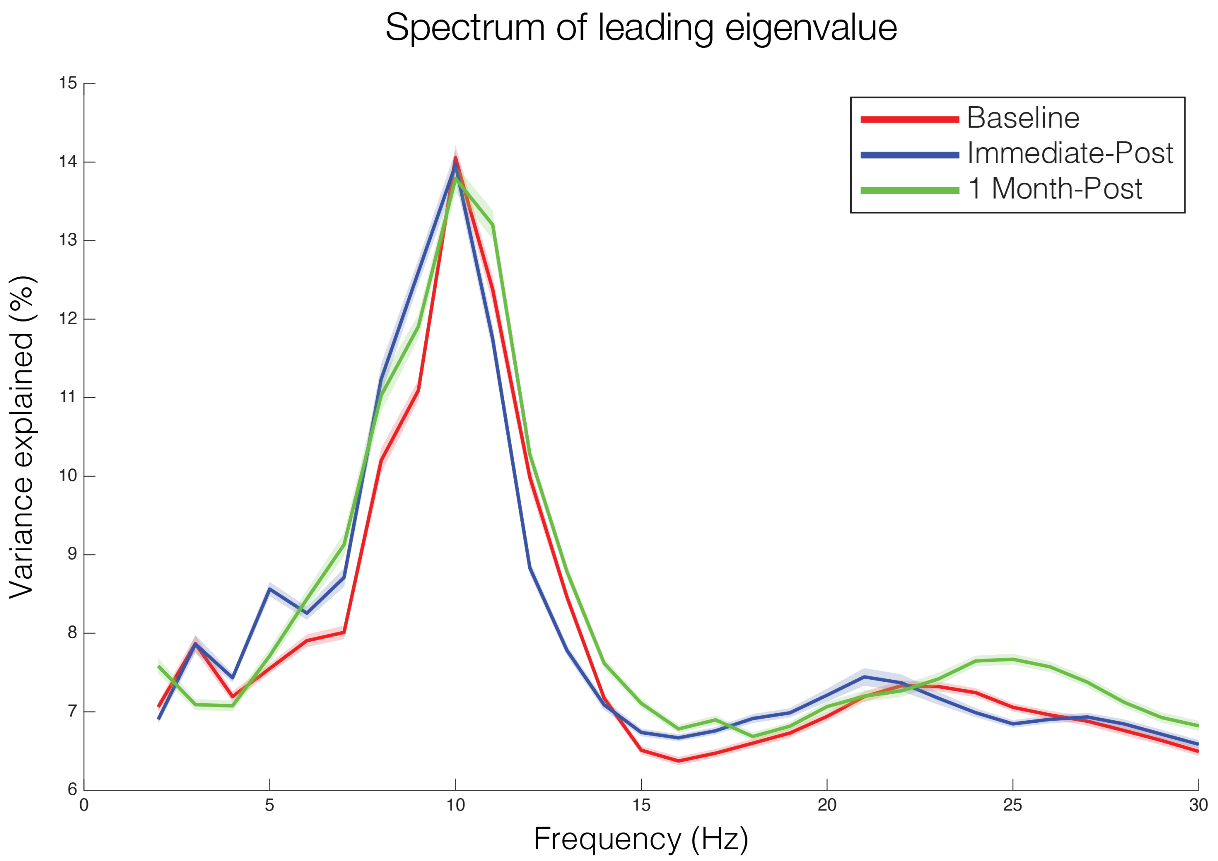


**Supplementary Figure 1. Ibogaine does not significantly affect the "prominence" or variance explained by any frequency-specific brain networks at any timepoints.** For each frequency, the leading eigenvalue is the greatest expression of narrow-band covariance, relative to broadband covariance, captured by any network. This corresponds to the prominence or strength of frequency-specific network organization. Like traditional EEG power spectra, the prominence peaks at 10 Hz. There were no significant differences between timepoints in the prominence of any frequency-specific brain networks. However, the decrease from baseline to immediate-post at 9 Hz, as well as the decrease from immediate-post to one month-post at 11 and 12 Hz, trended towards significance.

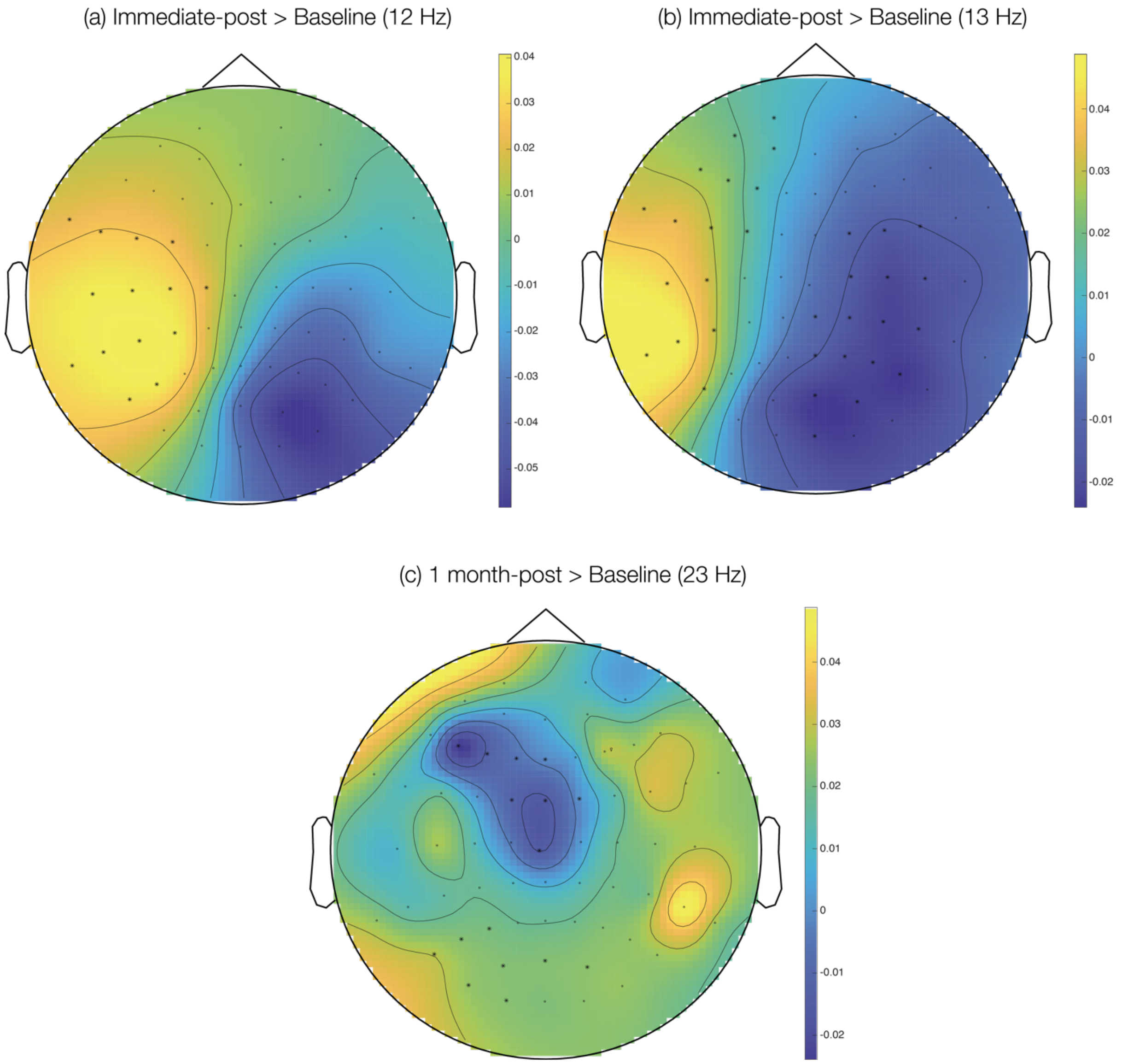


**Supplementary Figure 2. Ibogaine significantly alters the topography of high alpha and other high beta networks when an outlier participant is removed.** One subject exhibited aberrantly high network activations. When this subject was removed, changes in the topographies of three other networks became significant. (a, b) At 12 and 13 Hz, the topography became significantly more lateralized to the left hemisphere at immediate-post relative to baseline. (c) Shifts in the 23 Hz network at 1 month-post, relative to baseline, were similar to those of 24 Hz and 25 Hz, displaying a loss of activation at frontal left and central electrodes.

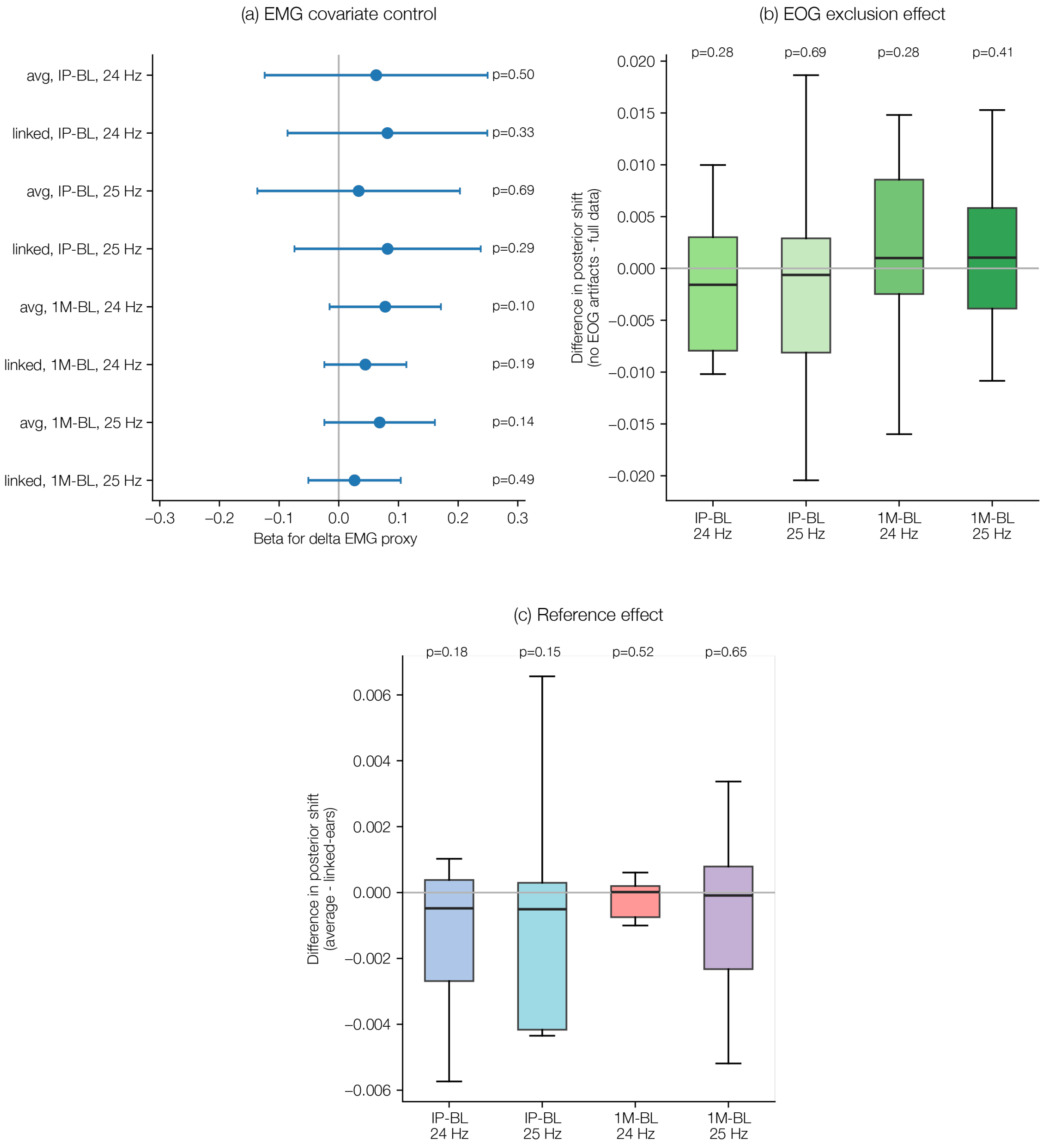


**Supplementary Figure 3. Control analyses of the high-beta posterior shift.** (a) EMG covariate control: Ordinary least squares estimates (β) for the association between change in EMG proxy (ΔEMG) and posterior shift, fit separately by timepoint (immediate-post vs. baseline, 1 month-post vs. baseline), frequency (24, 25 Hz), and reference scheme (average, linked-ears [see below]). Posterior shift was measured by subtracting the network activation patterns in the posterior electrodes from those of the anterior electrodes (**Supplementary Figure 9**). β was not significant for any of the comparisons. (b) EOG exclusion control: Boxplots of changes in posterior shift after excluding epochs with high EOG activity, as determined by proxy from the standard deviation of Fp1, Fp2, F7, and F8 activity. This exclusion had no significant effect on the posterior shift. (c) Reference robustness control: Boxplots of differences in posterior shift between reference schemes (average and linked-ears). The choice of reference did not significantly affect the posterior shift. IP = immediate-post; 1M = 1 month-post.

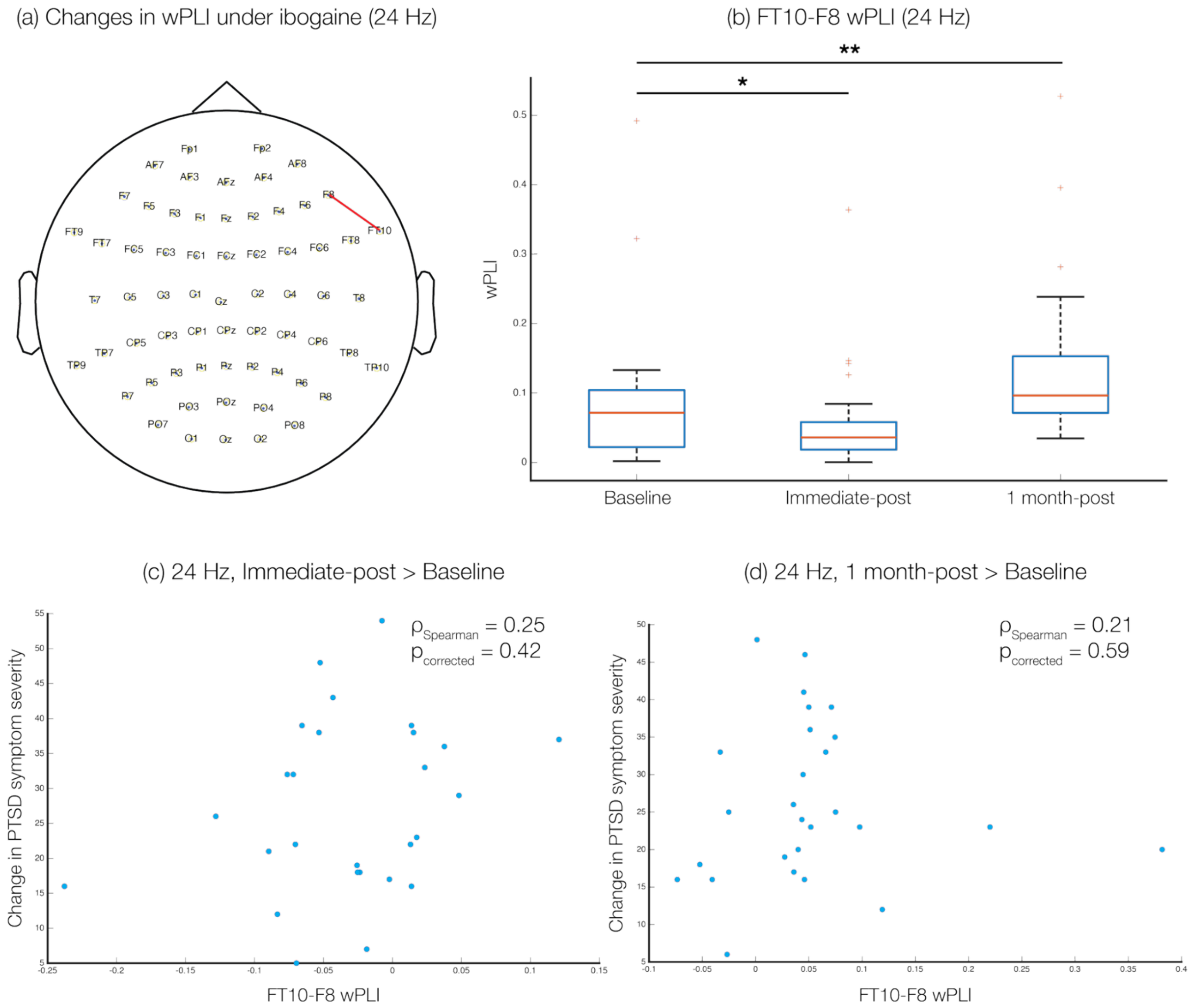


**Supplementary Figure 4. Traditional measures of functional connectivity like weighted phase lag index (wPLI) do not correlate with PTSD improvements.** (a) When network-based statistics are used to correct for multiple comparisons, there are no significant changes in wPLI under ibogaine. When false discovery rate (FDR) is used, ibogaine still does not have a significant effect on 25 Hz wPLI, but it significantly alters 24 Hz wPLI between only a single pair of electrodes, FT10 and F8. (b) Ibogaine significantly decreases 24 Hz FT10-F8 wPLI at the immediate-post timepoint and significantly increases it at the 1 month-post timepoint. (c, d) There is no significant correlation between PTSD improvements and FT10-F8 wPLI at either the immediate-post or 1 month-post timepoint.

(a) CAPS-5 B (Intrusion): Immediate-post > Baseline (24 Hz)

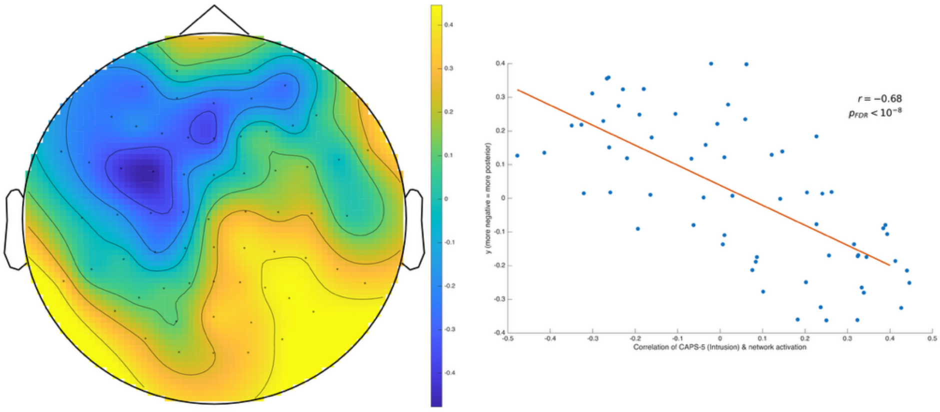


(b) CAPS-5 B (Intrusion): 1 month-post > Baseline (24 Hz)

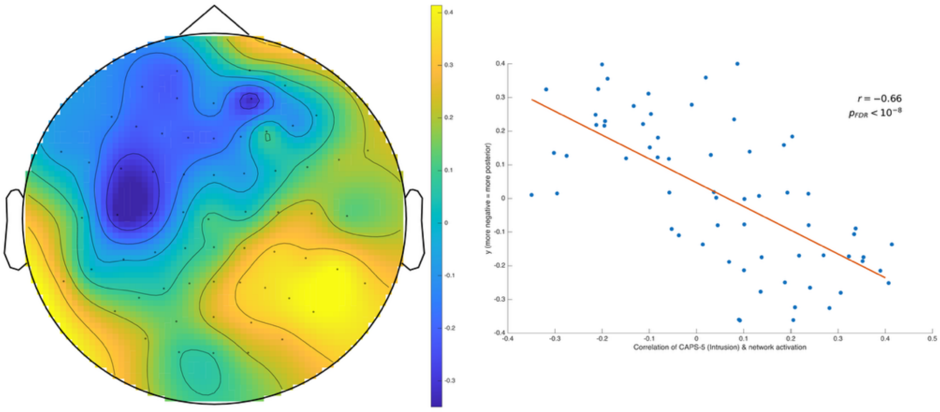


(c) CAPS-5 B (Intrusion): Immediate-post > Baseline (25 Hz)

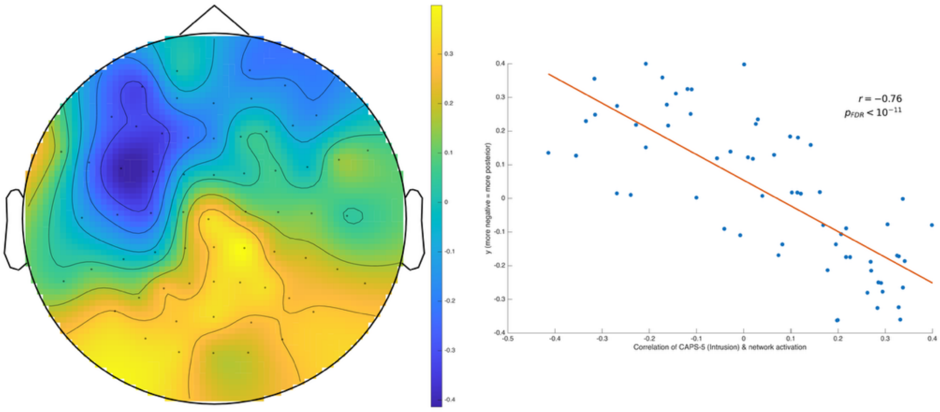


(d) CAPS-5 B (Intrusion): 1 month-post > Baseline (25 Hz)

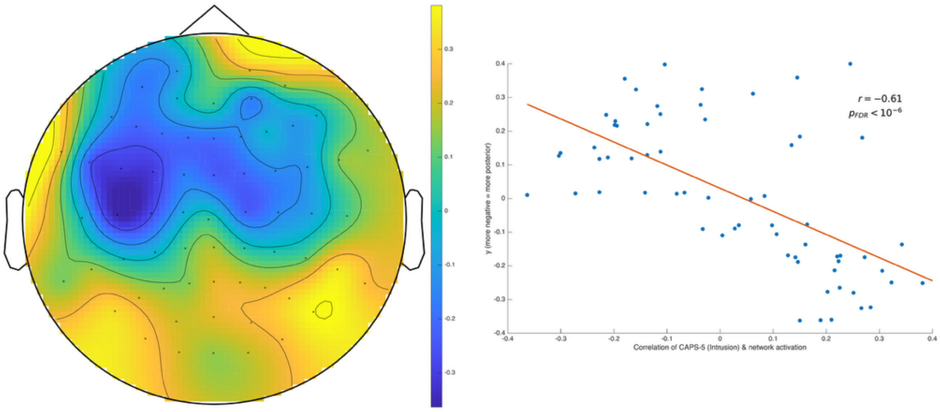

**Supplementary Figure 5. Posterior shifts in the high-beta network topography are significantly correlated with improvements in PTSD symptoms of intrusion, both immediately and one month after ibogaine.** Each electrode's network activation (the values shown in **Figure 2**) was correlated with changes in CAPS-5 subscale B scores, which reflect the intensity of intrusive traumatic memories and distress from trauma-related triggers. Changes in CAPS-B scores were multiplied by -1 such that increases in CAPS-B scores reflected improvement. With Pearson correlations, we then determined whether the strength of this association could be predicted by the *y*-coordinate of the electrode, i.e., its position along the anterior-posterior axis of the brain. At both 24 Hz and 25 Hz and at both immediate-post and one month-post, more posterior electrodes exhibited significantly more positive correlations between their network activation and improvements on CAPS-B, whereas more anterior electrodes exhibited significantly more negative correlations between their network activation and improvements on CAPS-B.

(a) CAPS-5 C (Avoidance): Immediate-post > Baseline (24 Hz)

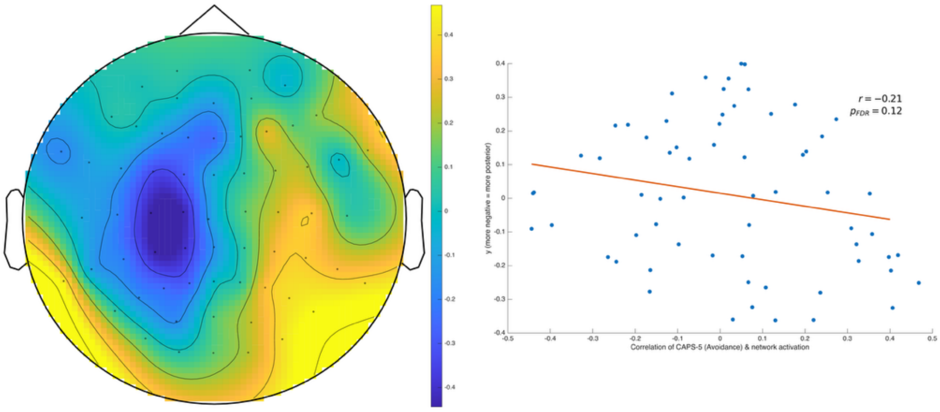


(b) CAPS-5 C (Avoidance): 1 month-post > Baseline (24 Hz)

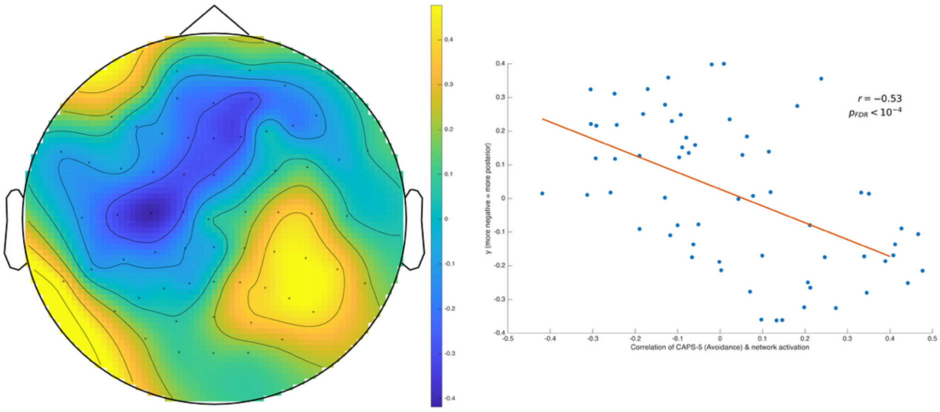


(c) CAPS-5 C (Avoidance): Immediate-post > Baseline (25 Hz)

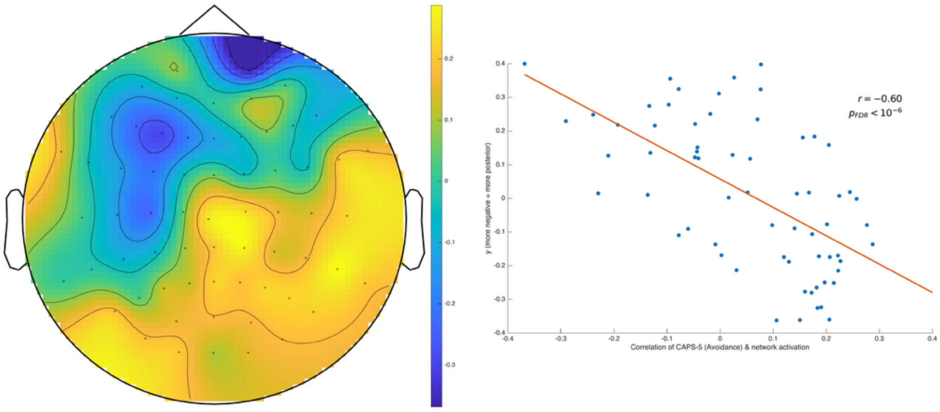


(d) CAPS-5 C (Avoidance): 1 month-post > Baseline (25 Hz)

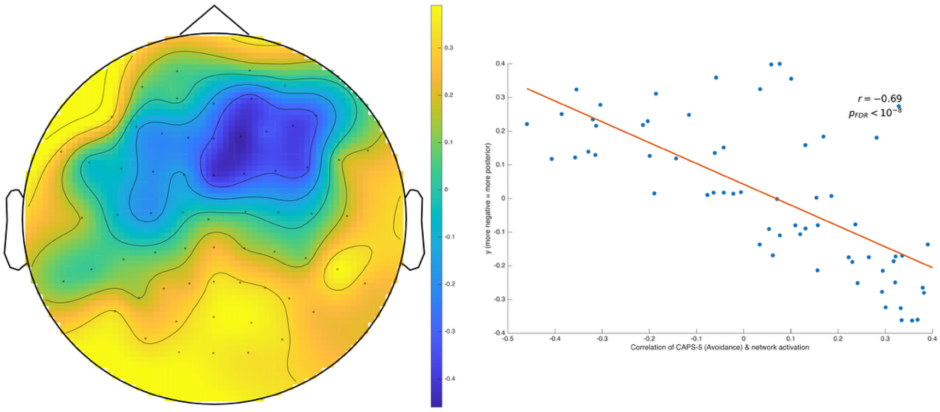

**Supplementary Figure 6. Posterior shifts in the high-beta network topography are significantly correlated with improvements in PTSD symptoms of avoidance.** Each electrode's network activation (the values shown in **Figure 2**) was correlated with changes in CAPS-5 subscale C scores, which reflect avoidance of trauma-related triggers, memories, thoughts, and feelings. Changes in CAPS-C scores were multiplied by -1 such that increases in CAPS-C scores reflected improvement. With Pearson correlations, we then determined whether the strength of this association could be predicted by the *y*-coordinate of the electrode, i.e., its position along the anterior-posterior axis of the brain. More posterior electrodes exhibited more positive correlations between their network activation and improvements on CAPS-C, whereas more anterior electrodes exhibited more negative correlations between their network activation and improvements on CAPS-C. This pattern reached statistical significance for the 25 Hz network at both immediate-post and one month-post, and for the 24 Hz network only at one month-post.

(a) CAPS-5 D (Cognition/Mood): Immediate-post > Baseline (24 Hz)

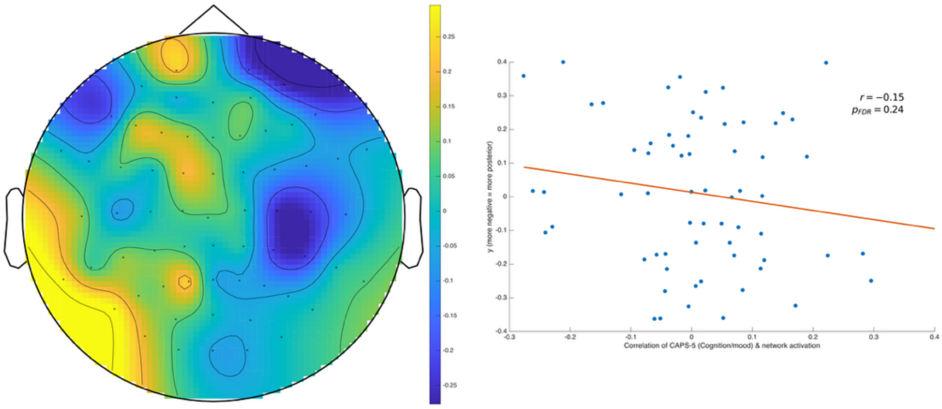


(b) CAPS-5 D (Cognition/Mood): 1 month-post > Baseline (24 Hz)

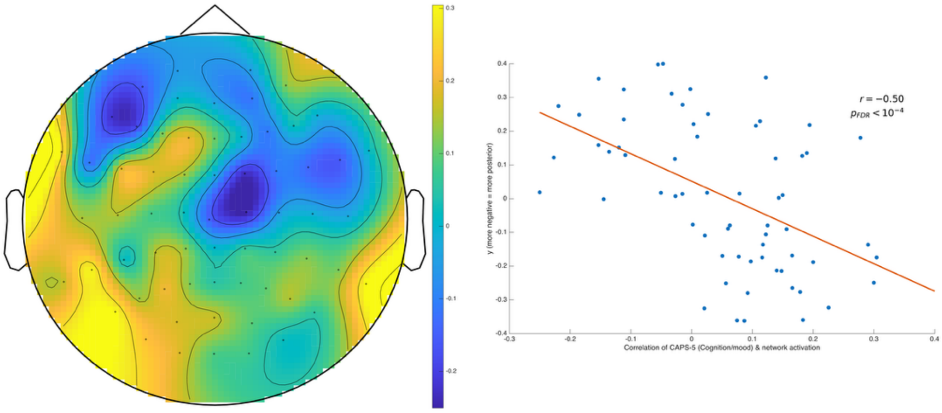


(c) CAPS-5 D (Cognition/Mood): Immediate-post > Baseline (25 Hz)

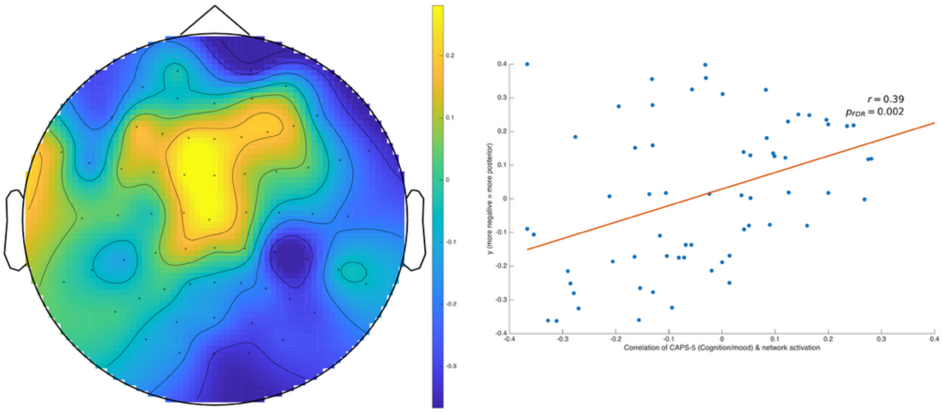


(d) CAPS-5 D (Cognition/Mood): 1 month-post > Baseline (25 Hz)

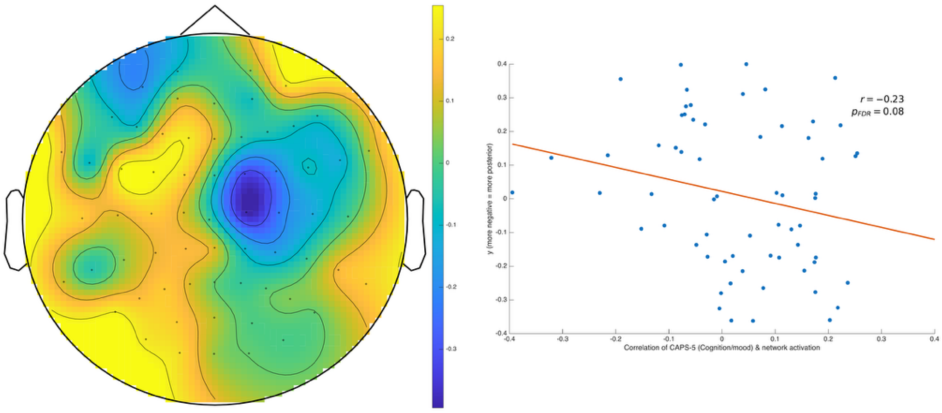

**Supplementary Figure 7. Posterior shifts in the high-beta network topography exhibit inconsistent correlations with improvements in PTSD symptoms of cognition and mood.** Each electrode's network activation (the values shown in **Figure 2**) was correlated with changes in CAPS-5 subscale D scores, which reflect impairments in memory of the trauma and negative alterations in feelings, thoughts, and beliefs. Changes in CAPS-D scores were multiplied by -1 such that increases in CAPS-D scores reflected improvement. With Pearson correlations, we then determined whether the strength of this association could be predicted by the *y*-coordinate of the electrode, i.e., its position along the anterior-posterior axis of the brain. For the 24 Hz network, more posterior electrodes exhibited significantly more positive correlations between their network activation and improvements on CAPS-D, whereas more anterior electrodes exhibited significantly more negative correlations between their network activation and improvements on CAPS-D; however, this pattern was only significant at the one month-post timepoint, not the immediate-post timepoint. The 25 Hz network displayed the opposite pattern; at the immediate-post timepoint (but not one month-post), the correlation between network activation and improvements on CAPS-D was more positive for more *anterior* electrodes.

(a) CAPS-5 E (Arousal/Reactivity): Immediate-post > Baseline (24 Hz)

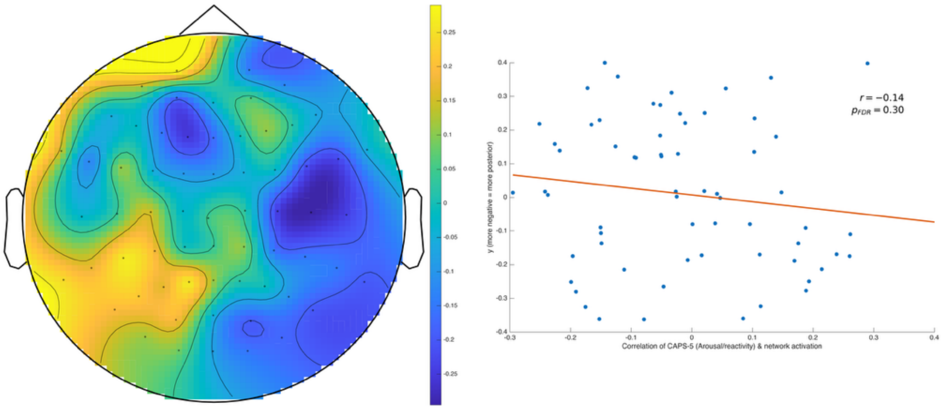


(b) CAPS-5 E (Arousal/Reactivity): 1 month-post > Baseline (24 Hz)

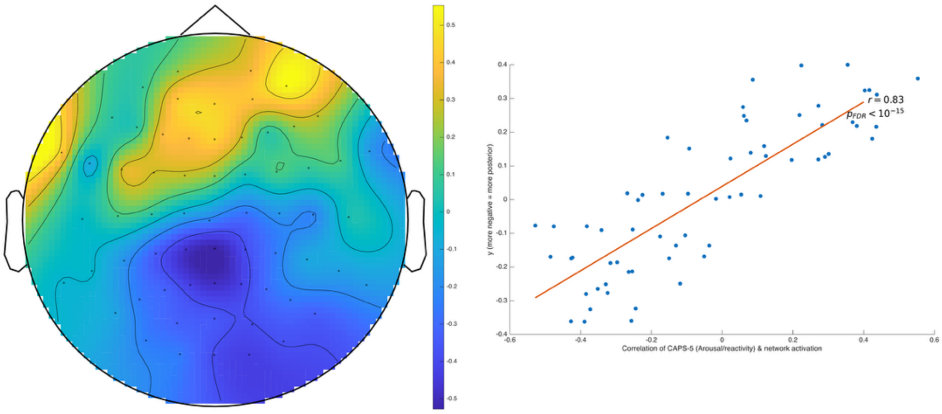


(c) CAPS-5 E (Arousal/Reactivity): Immediate-post > Baseline (25 Hz)

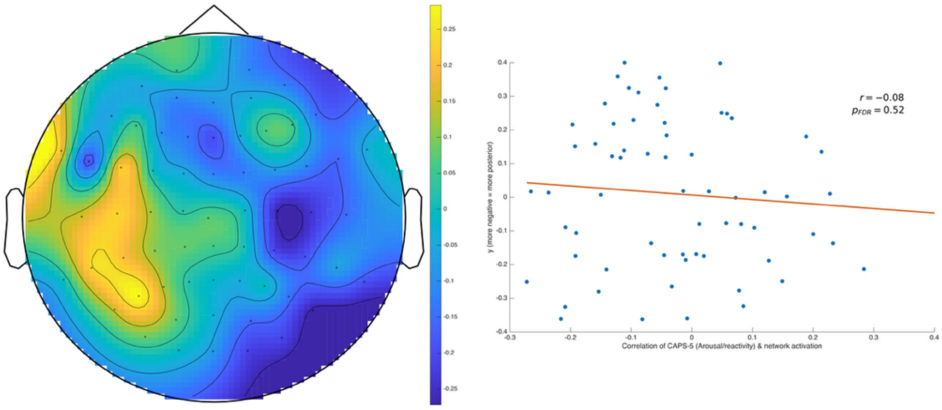


(d) CAPS-5 E (Arousal/Reactivity): 1 month-post > Baseline (25 Hz)

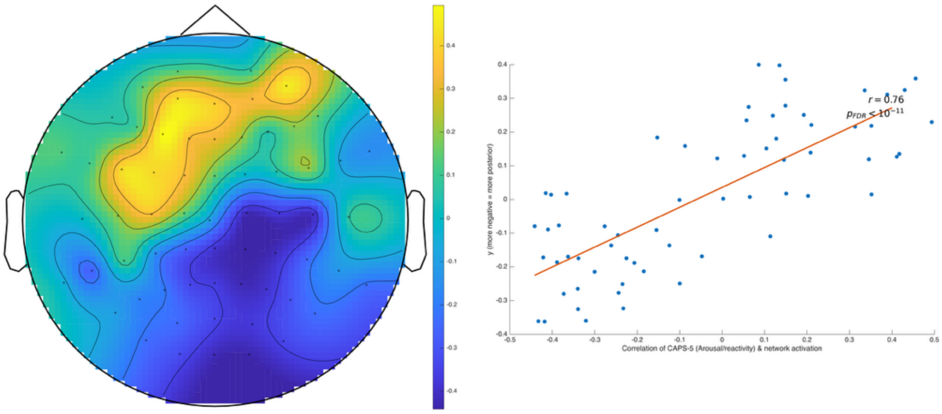

**Supplementary Figure 8. Posterior shifts in the high-beta network topography are anticorrelated with improvements in PTSD symptoms of arousal and reactivity.** Each electrode's network activation (the values shown in **Figure 2**) was correlated with changes in CAPS-5 subscale E scores, which reflect increases in arousal, reactivity, irritability, and hypervigilance. Changes in CAPS-E scores were multiplied by -1 such that increases in CAPS-E scores reflected improvement. With Pearson correlations, we then determined whether the strength of this association could be predicted by the *y*-coordinate of the electrode, i.e., its position along the anterior-posterior axis of the brain. For both the 24 and 25 Hz networks, more *anterior* electrodes exhibited significantly more positive correlations between their network activation and improvements on CAPS-E, whereas more *posterior* electrodes exhibited significantly more negative correlations between their network activation and improvements on CAPS-E. This pattern was observed only at the one month-post timepoint for each network, and contradicts the pattern of the CAPS-B and CAPS-C scores.

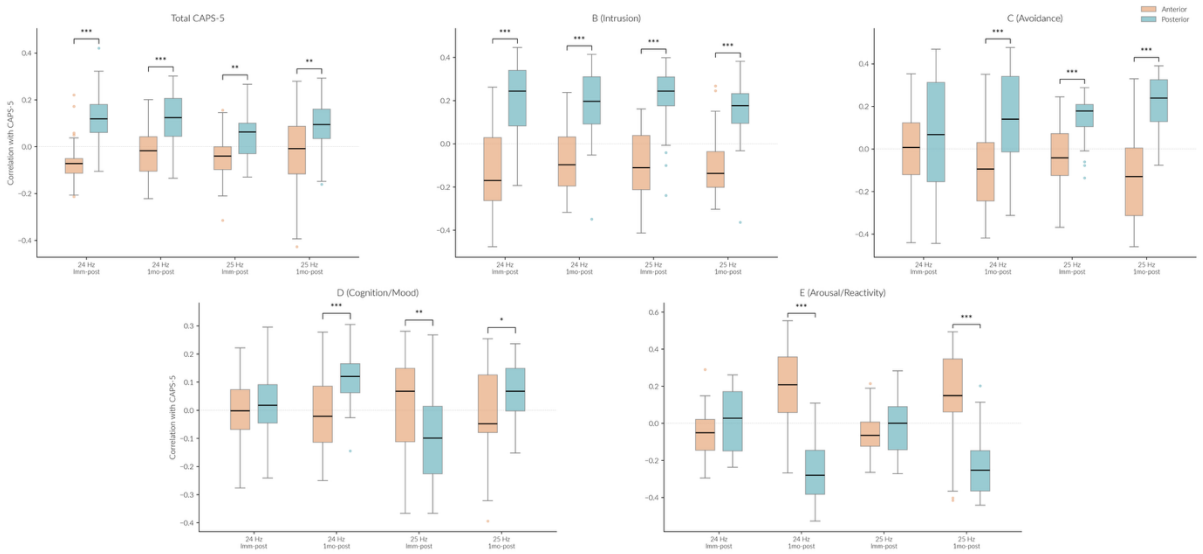


**Supplementary Figure 9. Different method of measuring posterior shift is consistent with previous results.** EEG topoplots denoting correlations between CAPS-5 scores and network activation (**Figure 4**, **Supplementary Figure 5-8**) were divided into anterior and posterior halves. Each subplot denotes whether there is a significant difference in correlation between the anterior (orange) and posterior (blue) halves at the immediate-post and one month-post timepoints (relative to baseline), for both the 24 and 25 Hz networks. Posterior electrodes generally exhibited more positive correlations than anterior electrodes, particularly for subscale B (Intrusion), where all four conditions were highly significant. Consistent with our previous results (**Supplementary Figure 8**), Criterion E (Arousal/Reactivity) showed a reversed pattern at the one-month timepoint, with anterior electrodes showing positive correlations and posterior electrodes showing negative correlations. *$p_{FDR} < 0.05$, **$p_{FDR} < 0.01$, ***$p_{FDR} < 0.001$.

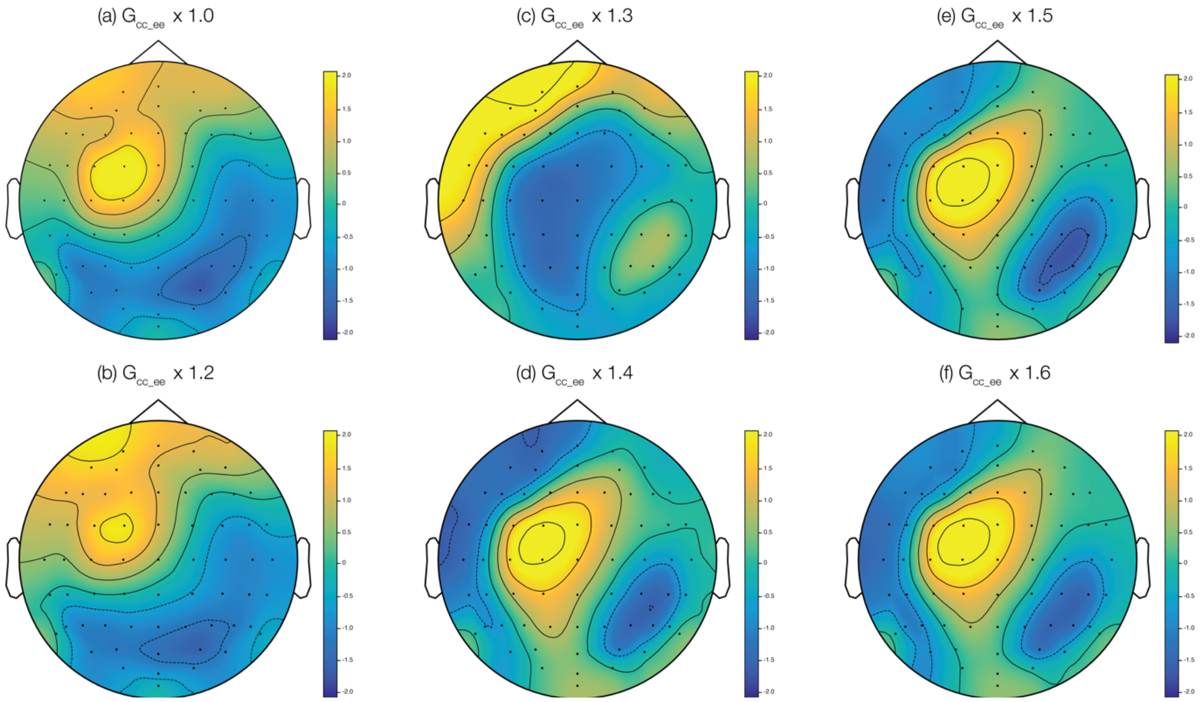


**Supplementary Figure 10. Increases in corticocortical gain, past a certain threshold, are also associated with decreases in frontal left activation in simulated high-beta networks.** (a) This is the same simulated 24 Hz network as shown in **Figure 5c**. (b, c) Elevating the gain of the connectivity from excitatory cortical to excitatory cortical neurons ($G_{cc\_ee}$) by 20% and 30%, respectively, increases the activation of frontal left electrodes in the simulated 24 Hz network. (d-f) However, increasing this gain by 40% or more leads to a drop in activation in these same electrodes.

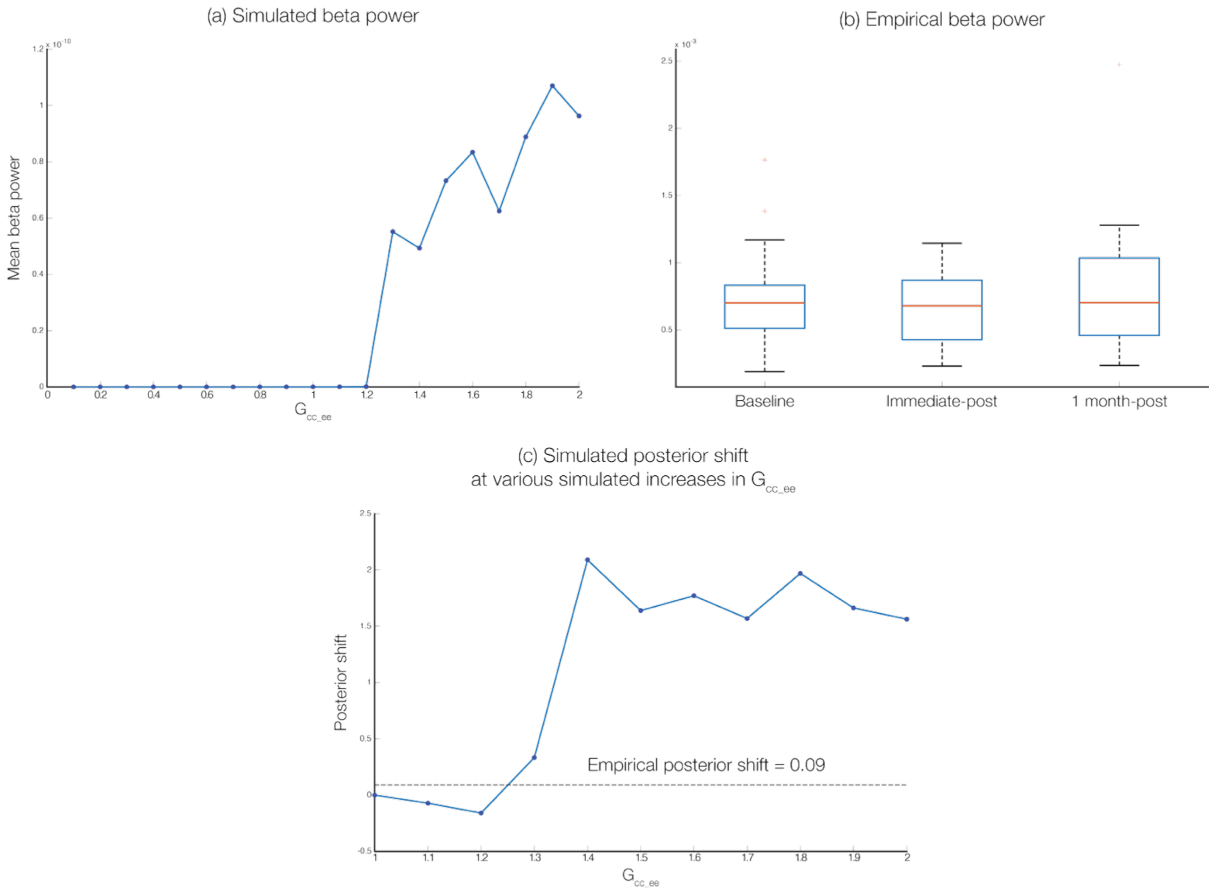


**Supplementary Figure 11. Increases in corticocortical gain are inconsistent with empirical beta power and leading eigenvalue of high-beta networks.** (a) When the gain of the connectivity between cortical excitatory and cortical excitatory ($G_{cc\_ee}$) is increased beyond 30%, simulated beta power, averaged across nodes, increases dramatically. Decreases in $G_{cc\_ee}$ do not substantially alter beta power. (b) Empirically, there is no significant change in beta power, averaged across electrodes, after ibogaine. (c) Although increases in $G_{cc\_ee}$ of 40% or greater are associated with a posterior shift, this simulated shift is much higher than the empirical shift.

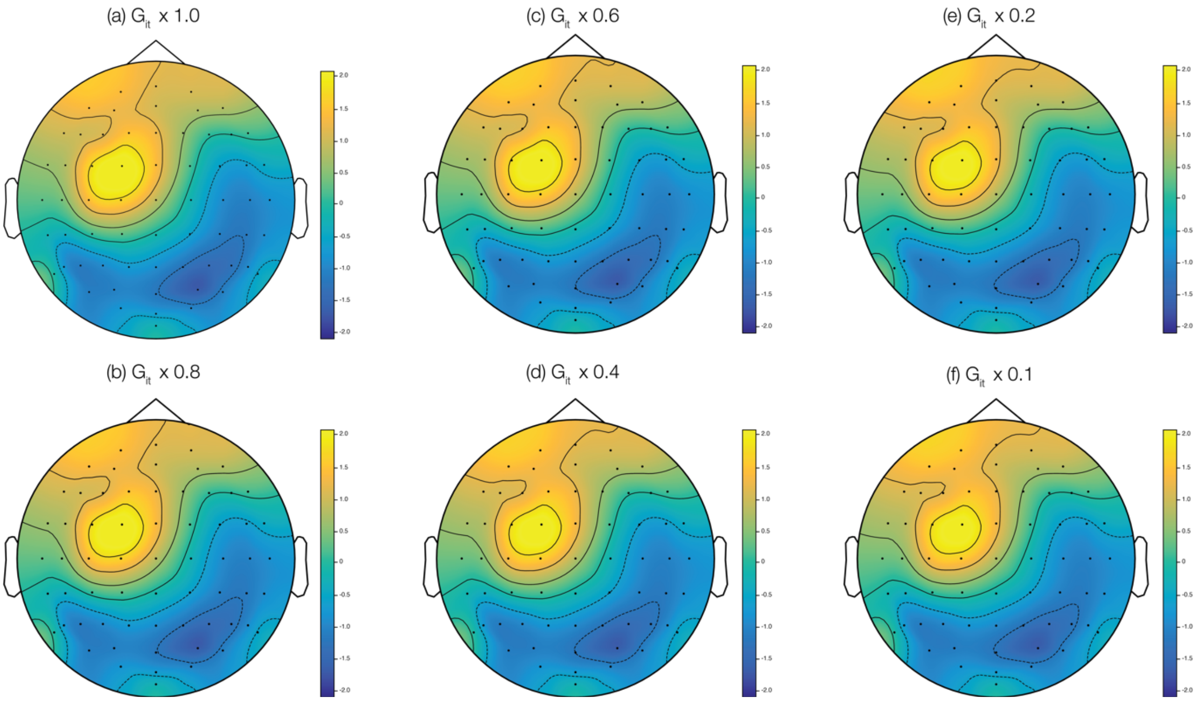


**Supplementary Figure 12. Decreases in intrathalamic gain do not lead to a loss of frontal left activation in simulated high-beta networks.** (a) This is the same simulated 24 Hz network as shown in **Figure 5c**. (b-f) Decreasing $G_{it}$, the gain of the connectivity between intrathalamic neurons (specifically, reticular thalamic and relay thalamic neurons), by 20%, 40%, 60%, 80%, and even 90%, respectively, does not cause a noticeable change in the topography of the 24 Hz network. Therefore, while ibogaine does significantly decrease intrathalamic gain at the immediate-post timepoint, this cannot explain the changes in the 24 Hz network topography.

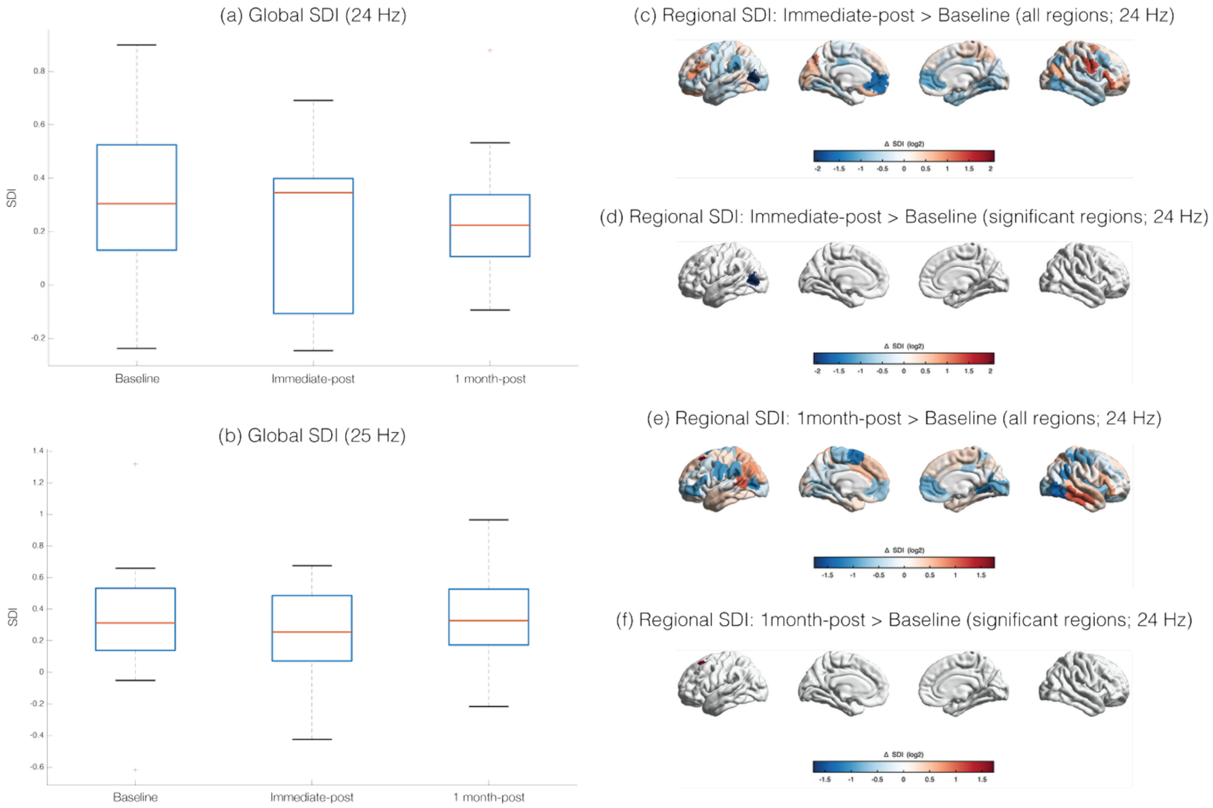


**Supplementary Figure 13. Ibogaine alters structure-function decoupling selectively in certain regions.** We measured the Structural-Decoupling Index (SDI) using structural connectomes, obtained from DWI data of the participants, and the regional network activation timeseries, obtained from applying FREQ-NESS to the source-reconstructed EEG data. (a-b) Ibogaine did not have a significant effect on global SDI, i.e., SDI averaged across regions, at either 24 or 25 Hz. (c-f) Ibogaine elicited widespread changes in regional SDI. However, the only significant changes were a decrease in SDI in the left inferior occipital cortex at the immediate-post timepoint and an increase in SDI in the left superior frontal gyrus at the one month-post timepoint.

| | Cluster Size | Max $t$ | Cluster $p$ | FDR-adjusted $p$ |
|---|---|---|---|---|
| 24 Hz, 12 days-post > Baseline | 1 | CP2: $t = 2.642$ | $p = 0.1084$ | $p_{corrected} = 0.1084$ |
| 25 Hz, 12 days-post > Baseline | 3 | CP2: $t = 3.176$ | $p = 0.0137$ | $p_{corrected} = 0.0273$ |

**Supplementary Table 1. Statistics of significant clusters of changes in high-beta network topographies in independent EEG dataset on ibogaine.** Cluster size, channel where the test statistic ($t$-value) reaches its absolute maximum and the value of that statistic, cluster-level $p$-value, and cluster-level $p$-value corrected across 2 frequencies (24 and 25 Hz).
